# mRNA structure dynamics identifies the embryonic RNA regulome

**DOI:** 10.1101/274290

**Authors:** Jean-Denis Beaudoin, Eva Maria Novoa, Charles E Vejnar, Valeria Yartseva, Carter Takacs, Manolis Kellis, Antonio J Giraldez

## Abstract

RNA folding plays a crucial role in RNA function. However, our knowledge of the global structure of the transcriptome is limited to steady-state conditions, hindering our understanding of how RNA structure dynamics influences gene function. Here, we have characterized mRNA structure dynamics during zebrafish development. We observe that on a global level, translation guides structure rather than structure guiding translation. We detect a decrease in structure in translated regions, and we identify the ribosome as a major remodeler of RNA structure *in vivo*. In contrast, we find that 3’-UTRs form highly folded structures *in vivo*, which can affect gene expression by modulating miRNA activity. Furthermore, we find that dynamic 3’-UTR structures encode RNA decay elements, including regulatory elements in *nanog* and *cyclin A1*, key maternal factors orchestrating the maternal-to-zygotic transition. These results reveal a central role of RNA structure dynamics in gene regulatory programs.

## Introduction

RNA carries out a broad range of functions,^1^ and RNA structure has emerged as a fundamental regulatory mechanism, modulating various post-transcriptional events such as splicing^2^, subcellular localization^3^, translation^4,5^ and decay^6^. RNA probing reagents combined with high-throughput sequencing^7^ allow the interrogation of RNA structure in a transcriptome-wide manner *in vitro* and *in vivo*^8,9^. These approaches have identified common features in RNA structures and their regulation at steady-state^10–14^, suggesting that RNAs tend to be unfolded *in vivo*^13,15^. However, the cellular factors that remodel RNA folding *in vivo* remain unknown.

While the ribosome possesses a constitutive mRNA helicase activity^16^, stable RNA structures found within coding regions can reduce the rate of translation *in vitro*^17,18^. Similarly, the intrinsic structure adopted by coding regions instruct the rate of translation in bacteria, where sequences forming stable structures show reduced translation *in vivo*^19^. In addition, stable structures located in 5’UTR or surrounding the AUG initiation codon repress translation and modulate protein output in bacteria and eukaryotes^20–23^. However, in plant and yeast, highly structured mRNA *in vitro* correlated with higher ribosome density and protein output^9,24^, yet, other studies found that RNA structure in yeast were not correlated with translation efficiency^13^.

Previous studies have focused on the global analysis of RNA structures in cells at steady-state. However, in an organism, critical developmental and metabolic decisions are often made when cells transition between cellular states. Consequently, to understand the relationship between RNA structure, regulation, and function *in vivo*, it is paramount to determine how RNA structures change in dynamic cellular environments. During the maternal-to-zygotic transition (MZT) in animals, the embryo is initially transcriptionally silent, and gene expression is primarily orchestrated by post-transcriptional regulation of maternally-deposited mRNA translation and decay^25,26^. These regulatory pathways are central to embryogenesis, controlling mRNA translation, the cell cycle, and the activation of the zygotic genome^25,27^. Among the set of maternally-deposited mRNAs, the transcription factors *nanog, oct4*, and *sox19b* are highly translated. Their combined function is required to activate a large fraction of the zygotic program^28,29^. One of the first zygotically transcribed RNAs is the microRNA miR-430^28,30^, which causes translation repression, deadenylation and clearance of hundreds of maternal mRNAs during the MZT^31–33^. Thus, the MZT provides an ideal system to understand how the entire cellular population of RNA structures is remodeled across changing cellular states, and how these changes may impact gene regulation. Here, we analyze mRNA structural dynamics during zebrafish embryogenesis to investigate the relationship between RNA structure and gene regulation *in vivo*.

## Results

### Dynamic structure reflects translation changes

We analyzed the structure of the zebrafish transcriptome during the MZT using DMS-seq (Fig 1a), which we validated as an accurate readout of *in vivo* RNA structure (Extended Data Fig. 1). Our analysis revealed global changes in mRNA structure during the MZT (Extended Data Fig. 1i-k). Changes in translation, analyzed by ribosome footprinting^34^, were correlated with global mRNA accessibility changes in coding sequences (CDS) and 5’-UTRs, but not in 3’-UTRs (Fig. 1b and Extended Data Fig. 1l). We analyzed differentially structured regions over development as 100-nt sliding windows (Fig. 1c) and observed that mRNAs with decreasing rates of translation between 2 and 6 hours post-fertilization (hpf) increase in structure (orange), and those whose translation increases become less structured (turquoise) (Fig. 1d). These results indicate that translation and mRNA structure are anti-correlated *in vivo*, in contrast to previous reports^9,13,24,35^. Moreover, we compared the accessibilities of the different mRNA regions (5’-UTR, CDS and 3’-UTR) and found that highly translated mRNAs display a greater accessibility in their CDS (*P*=2.3e-118) and, to a lesser extent, in their 5’-UTRs (*P*=1.9e-11) (Fig. 1e and Extended Data Fig. 2a).

**Figure 1.**
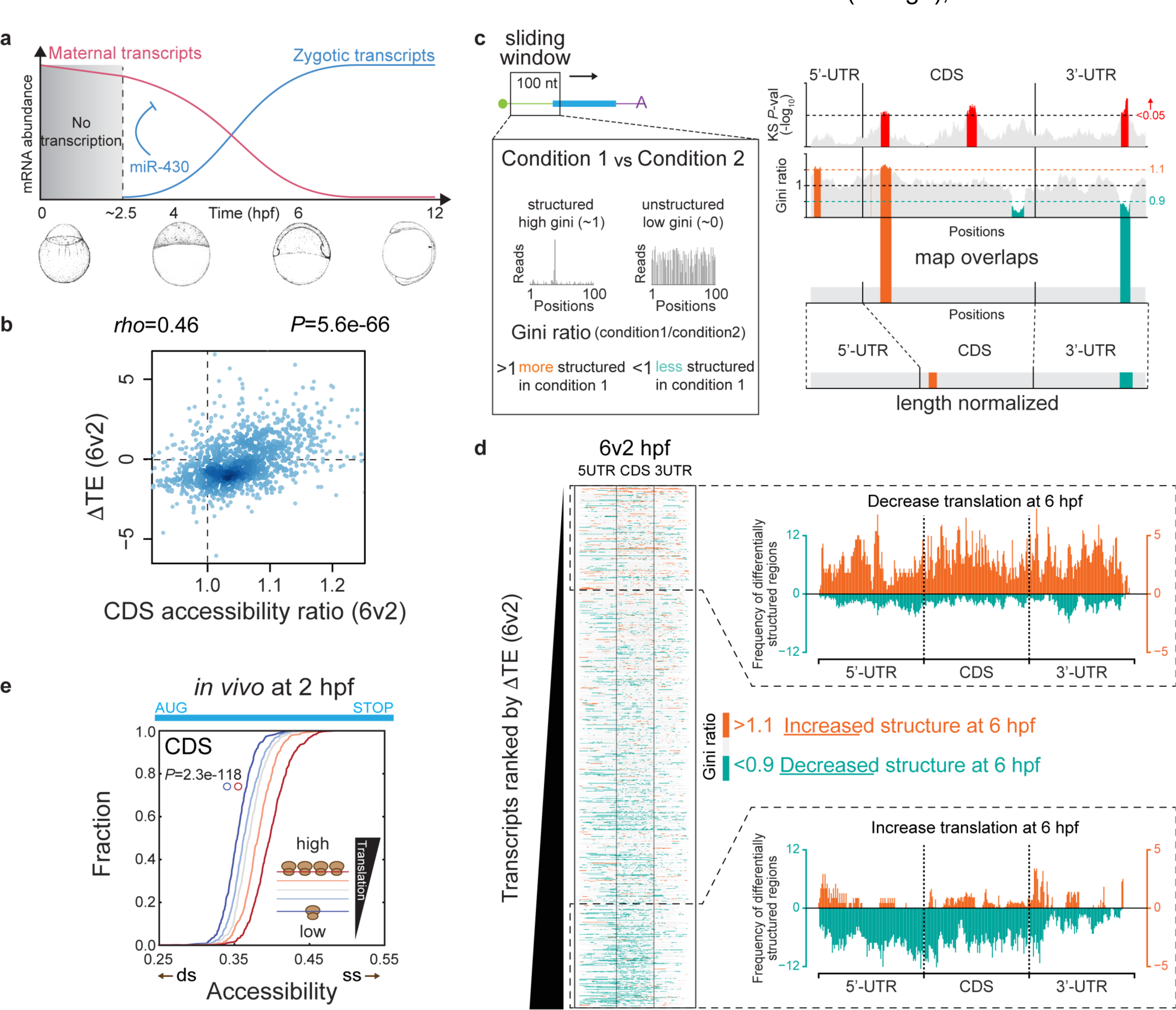
Relationship between mRNA structure and translation during the MZT. **a**, Schematic view of the profound transcriptomic remodeling that occurs during the maternal-to-zygotic transition. After a transcriptionally silent period (gray), the maternal program (pink) transitions to a zygotic program (blue). **b**, Correlation between the per-transcript changes in translation efficiency (6-2 hpf) and CDS accessibility (6/2 hpf). **c**, Methodology used to identify local RNA structure changes between conditions. Differentially structured (DS) sliding windows are defined as those that are significantly changing (*P*<0.05) based on the Kolmogorov-Smirnov test, and show an increase (>1.1) or decrease (<0.9) in their Gini index ratio. **d**, In the left panel, structure changes of each 100-nt window (Gini ratio 6/2 hpf) along each transcript (total of 1,273) are shown. Transcripts are ranked by their changes in translation efficiency during the MZT (6-2 hpf). Each transcript region (5’-UTR, CDS, 3’-UTR) has been normalized by its length. In the right panel, cumulative distributions of differentially structured windows for transcripts that increase (top 20%, upper panel) and decrease (bottom 20%, lower panel) translation are shown. **e**, Cumulative distribution of global CDS accessibilities *in vivo*, showing that highly translated mRNAs exhibit increased CDS accessibility *in vivo* at 2 hpf, a stage at which the rate of translation spans ∼10,000-fold between highly and poorly translated mRNAs. Transcripts have been binned into quintiles according to their translation efficiency. *P*-value was computed using a one-sided Mann-Whitney *U* test.

**Figure 2.**
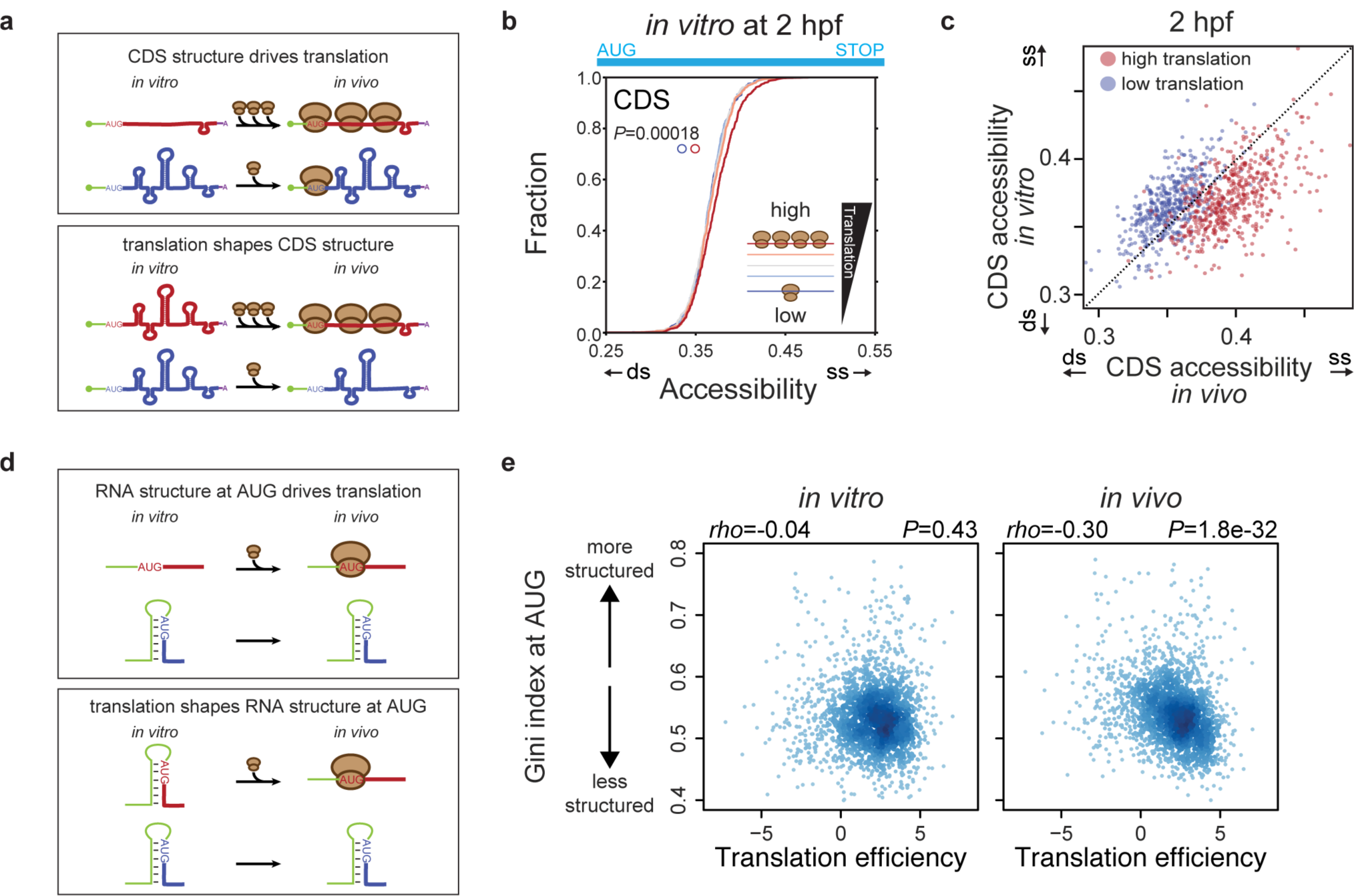
Globally, the mRNA intrinsic structure is not a main driver of translation. **a**, Schematic representation of the two models that can explain the observed anti-correlation between translation efficiency and RNA structure in CDS regions *in vivo*. In the first model, pre-existing CDS RNA structures would drive translation (discarded by our analysis, upper panel). In a second model, translating ribosomes would be responsible for the unfolding of CDS RNA structures, (consistent with our analysis, lower panel). **b**, Cumulative distributions of global CDS RNA accessibilities *in vitro* at 2 hpf, displaying no difference among mRNAs with different translation efficiency. Transcripts have been binned into quintiles according to their translation efficiency. *P*-value was computed a using one-sided Mann-Whitney *U* test. **c**, Scatterplot between *in vivo* and *in vitro* CDS accessibilities at 2 hpf. Note higher accessibility *in vivo* for highly translated mRNAs (red dots) and lower accessibility *in vivo* for poorly translated mRNAs (blue dots). **d**, Schematic view of the two models where RNA structure at the AUG initiation codon is either the cause (top) or the result (bottom) of mRNA translation. **e**, Correlation between translation efficiency and the structure at AUG regions, for both *in vitro* (left) and *in vivo* (right) conditions.

Pre-existing RNA structure differences might determine mRNA translation, where less structured mRNAs would be more accessible and, consequently, translated more effectively (Fig. 2a, upper panel). Alternatively, high translation rates might lead to lower structure *in vivo* due to constant mRNA unfolding by the ribosome (Fig. 2a, lower panel). To distinguish between these scenarios, we compared the accessibility of transcripts analyzed *in vitro*, binned by their translation efficiency *in vivo* (see Methods) (Fig. 2b and Extended Data Fig. 2a). We found that lowly and highly translated mRNAs show similar CDS accessibility *in vitro*, distinct from *in vivo* (Fig. 2b, c and Extended Data Fig. 2a-c), suggesting that the differences in mRNA accessibility between highly and lowly translated mRNAs are not intrinsic to the nucleotide sequence. While very stable structures in the 5’-UTR clearly disrupt translation^20,22,23^, we observed no correlation between translation and structure of 5’-UTRs or the AUG initiation codons *in vitro* (Fig. 2d, e and Extended Data Fig. 2d-g). Thus, on a global level, RNA structure in 5’-UTR and CDS regions is not a major determinant of translation across the transcriptome in a vertebrate embryo.

### Ribosomes shape mRNA structures

The helicase activity of the ribosome unfolds RNAs *in vitro* during translocation^16–18^. However, in yeast there is no correlation between RNA structure and translation efficiency (TE)^13^, suggesting that ribosome activity does not account for the differences between mRNA structures *in vivo* and *in vitro.* To determine the role of ribosomes in structural remodeling *in vivo*, we examined RNA structural changes in embryos incubated with inhibitors of translation initiation (pateamine A; PatA)^36,37^ or elongation (cycloheximide; CHX)^38^. Ribosome profiling of 64-cell embryos (2 hpf) treated with PatA revealed a prominent decrease (∼34-fold) in translation efficiency compared to untreated embryos (Fig. 3a, b). mRNAs most sensitive to PatA treatment had longer 5’-UTRs (*P*=2.1e-42), consistent with a loss of the PatA target eIF4A during scanning by the pre-initiation complex^36,37^ (Fig. 3a, c). In contrast, mitochondrial transcripts showed no change in translation (Fig. 3d), consistent with the observation that their translation is independent of eIF4A^39^. In PatA-treated embryos, we found an overall decrease in accessibility of the CDS regions of highly translated mRNAs (*P*=5.3e-35) (Fig. 3e and Extended Data Fig. 3a). Similarly, regions with higher ribosome footprint densities form more stable RNA conformations after PatA treatment, similar to those observed *in vitro* (Fig. 3f and Extended Data Fig. 3b, c). In contrast to PatA-treated embryos, CHX treatment did not decrease average CDS accessibility (Extended Data Fig. 3d), suggesting that even when the ribosomes are stalled^38^, the residency of the ribosome on the mRNA during translation plays a major role in remodeling mRNA structure. Therefore, translation initiation and ribosomal entry are needed to locally favor the formation of less stable alternative RNA structures.

**Figure 3.**
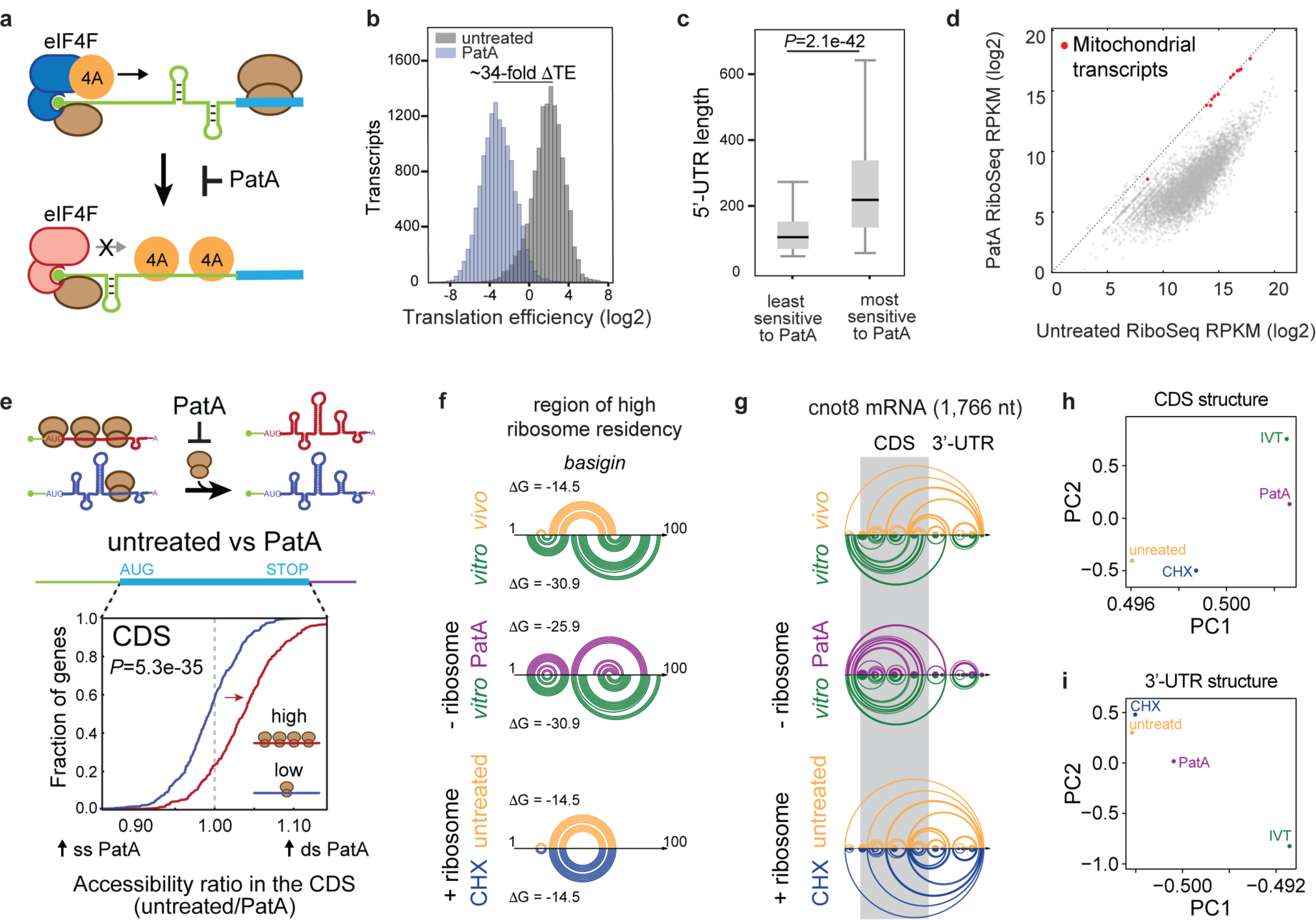
Ribosomes promote alternative mRNA structures in the early embryo. **a**, Schematic view of the PatA translation inhibitory mechanism. PatA reduces the levels of functional eIF4F initiation complex, which is essential for cap-dependent translation, by trapping eIF4A onto mRNAs and ectopically enhancing its RNA helicase activity. **b**, Distribution of translation efficiencies (TE) of PatA-treated (blue) and untreated (gray) embryos. **c**, Correlation between 5’-UTR length and sensitivity to PatA treatment. The least and most sensitive to PatA mRNA subgroups correspond to the top and bottom quintiles of ΔTE (PatA-untreated), respectively. *P*-value was computed using a one-sided Mann-Whitney *U* test. Box spans first to last quartiles and whiskers represent 1.5× the interquartile range. **d**, Scatter plot of the per-transcript ribosome footprints in PatA-treated versus untreated embryos. Mitochondrial genes (red), which are unaffected by PatA treatment, remain unchanged. **e**, Schematic representation depicting the impact of PatA, along with its corresponding cumulative distribution of CDS accessibility ratios (untreated/PatA). Note the decrease in accessibility (red arrow) in highly translated mRNAs in PatA compared to untreated. *P*-value was computed using a one-sided Mann-Whitney *U* test. **f**, Arc plots of DMS-seq guided RNA secondary structure predictions of a high ribosome occupancy region found in the *basigin* gene, for *in vivo* (yellow) *in vitro* (green), untreated (yellow), PatA-treated (purple) or CHX-treated (blue) samples. Each arc represents a base pair interaction. **g**, Arc plots of DMS-seq guided RNA secondary structure predictions of the full-length *cnot8* mRNA using SeqFold. The CDS region has been shadowed in gray. **h**, **i**, Principal Component Analysis biplots of the SeqFold mRNA structure predictions of the 4 tested conditions (untreated, *in vitro*, PatA-treated and CHX-treated), of CDS (**h**) and 3’-UTR (**i**) regions.

To evaluate the impact of the ribosome remodeling activity on mRNA structures, we compared the RNA conformations adopted by full-length transcripts in each probing condition using SeqFold^40^. We found that the same transcript could favor different conformations depending on its translation status (Fig. 3g and Extended Data Fig. 3e). Principal component analysis (PCA) on the structural data from 1,143 transcripts (see Methods) revealed that in absence of ribosomes (*in vitro* or in the presence of PatA), individual mRNAs exhibit similar CDS and 3’-UTR structures, distinct from those formed in the presence of ribosomes (untreated and CHX) (Fig. 3h, i). Finally, translation of upstream open reading frames (uORF) located in 5’-UTRs leads to a restructuring of 5’-UTR structures *in vivo* (Extended Data Fig. 4). Thus, the ribosome increases mRNA accessibility, and promotes alternative RNA conformations throughout the mRNA.

**Figure 4.**
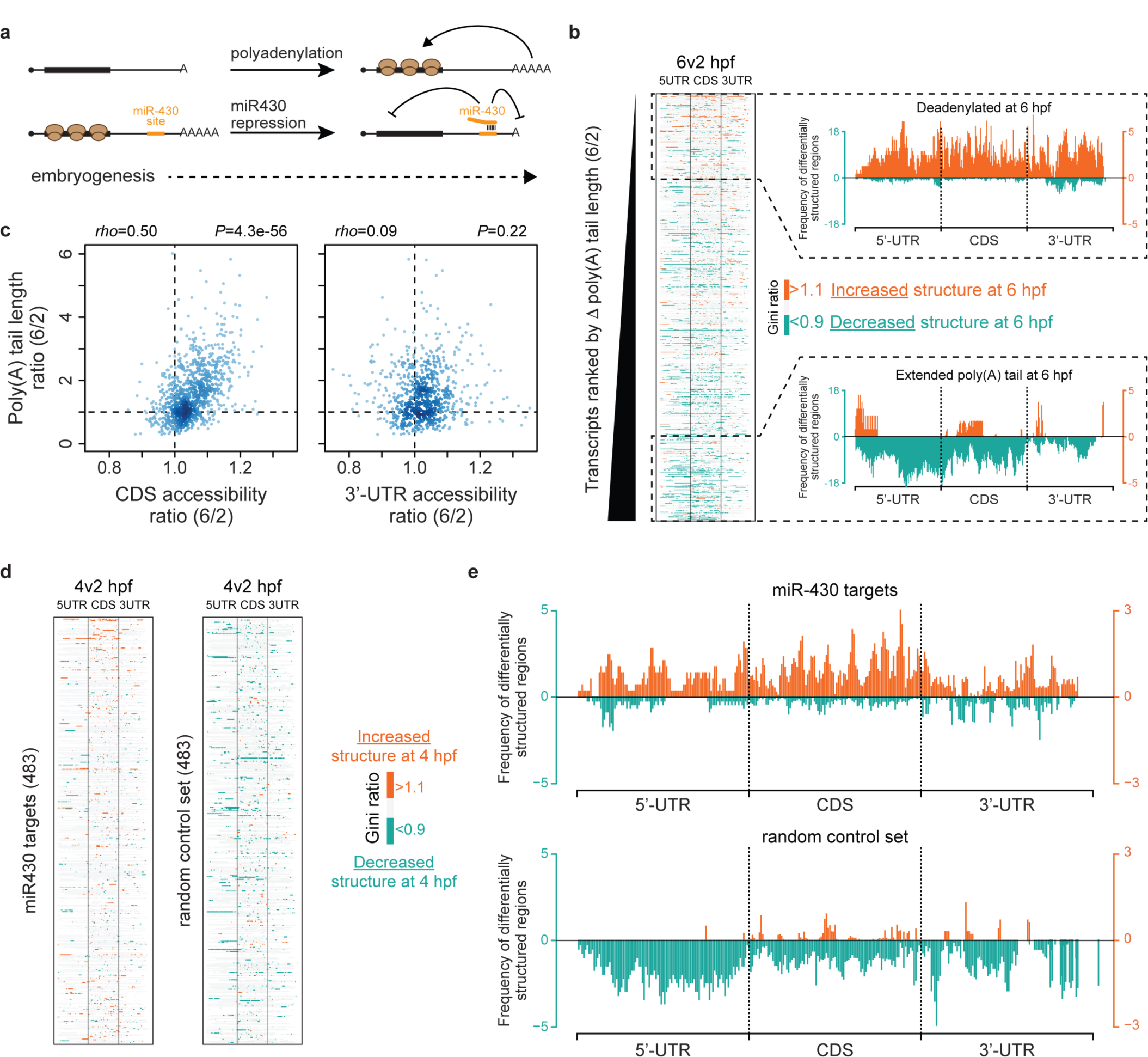
Polyadenylation and miR-430 activity influence mRNA structures. **a**, Schematic view of the impact of poly(A) tail length changes and miRNA-mediated repression on translation during early embryogenesis. **b**, In the left panel, the structure changes along each transcript (total of 965 mRNAs with sufficient coverage in DMS-seq and poly(A) tail length profiling experiments at both 2 and 6 hpf), ranked by their changes in poly(A) tail length (6/2 hpf). Structure changes have been identified by computing the Gini ratio for each 100-nt sliding window (6/2 hpf). In the right panel, the cumulative distribution of differentially structured windows along the transcripts, for both the top 20% (upper panel) and bottom 20% (lower panel) transcripts, binned by their poly(A) tail length changes, are shown. **c**,Correlation between changes in poly(A) tail length and accessibility (6/2 hpf) is shown, for both CDS and 3’-UTR regions. **d**, mRNA structure changes along each of the miR-430 targets (483) or a random set of transcripts (483). Regions with structure changes have been identified by computing the Gini ratio for each 100-nt sliding window (4/2 hpf). **e**, Metaplot of the differentially structured regions along the transcripts, comparing 2 and 4 hpf (4v2), for both miR-430 targets and randomly chosen transcripts.

### Translation-dependent mRNA structure

We then evaluated translation control mechanisms during the MZT for their impact on mRNA structure regulation (Fig. 4a). Polyadenylation increases translation efficiency during early embryogenesis^33,41^ (Fig. 4a). We found that mRNAs with longer poly(A) tails show higher accessibilities in the CDS at 2 and 4 hpf (*rho* of 0.42 and 0.36, respectively), but not in the 3’-UTR (Extended Data Fig. 5a). This effect was reduced following PatA treatment, and lost *in vitro* (*rho* of 0.29 and 0.01, respectively) (Extended Data Fig. 5b), consistent with a translation effect specific to the CDS. Moreover, mRNAs with extended poly(A) tails during the MZT displayed a decrease in structure, while deadenylated transcripts showed an increased structure (Fig. 4b, c). These results suggest that changes in polyadenylation during development influence mRNA structure, an effect that depends on the translation of the mRNA.

**Figure 5.**
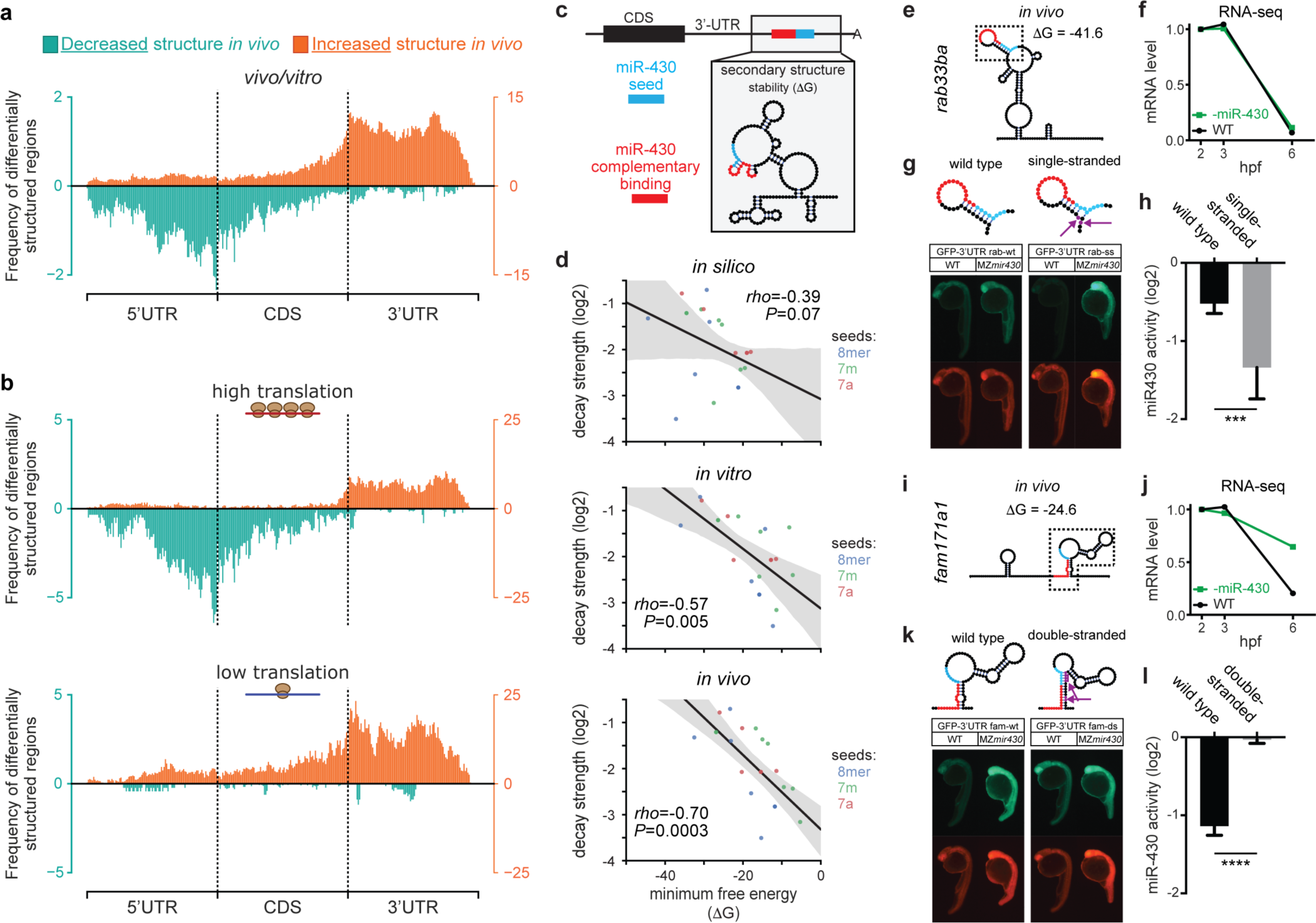
3’-UTR regions are more structured in the cell than *in vitro*, which influence gene expression. **a**, Distribution of differentially structured windows along the transcripts, comparing *in vivo* and *in vitro* conditions (*vivo*/*vitro*). Regions with increased structure *in vivo* are depicted in orange, whereas those with decreased structure *in vivo* are shown in turquoise. **b**, Distribution of differentially structured windows, comparing *in vivo* and *in vitro* conditions, for transcripts that have either high translation efficiency (top 20%, upper panel) or low translation efficiency (bottom 20%, lower panel). **c**, Cartoon representation of the RNA secondary structure of the region centered on a miR-430 target site, highlighting regions complementary to the miR-430 seed (cyan) and its complementary binding region within the 3’-UTR (red). **d**, Correlation between the decay strength and the stability of the RNA structure of 18 endogenous miR-430 binding sites,where the RNA secondary structure has been predicted using either an *in silico* approach (top), DMS-seq guided *in vitro* structural data (middle) or DMS-Seq guided *in vivo* structural data (bottom). **e**, **i**, *In vivo* predicted secondary structure and stability (ΔG) of the 200-nt region centered on the miR430 target site found in the *rab33ba* 3’-UTR (**e**) and in the *fam171a1* 3’-UTR (**i**). **f**, **j**, Changes in mRNA abundance during the MZT of the endogenous *rab33ba* (**f**) and *fam171a1* (**j**) transcripts in wild type conditions (black) or in conditions where miR-430 activity is inhibited by a tinyLNA complementary to the miR-430 seed (green). **g**, **k**, Fluorescence microscopy of GFP reporters at 24 hpf harboring either a wild type sequence of the rab33ba (rab-wt) or the fam171a1 (fam-wt) target sites, compared to a destabilized version of the rab33ba (rab33ba single-stranded, rab-ss) or a stabilized version of the fam171a1 (fam171a1 double-stranded, fam-ds) target sites in their 3’-UTR. The specific mutations of the rab-ss and fam-ds constructs, responsible for either disrupting or extending a stem, are highlighted in purple and with arrows. DsRed mRNA was co-injected as control. Reporters were injected in both wild type (WT) and in maternal-zygotic mutant of miR-430 (*MZmir430*) lacking the whole miR-430 locus. **h**, **l**, MiR-430 activity calculated from the fluorescence of the GFP normalized by the DsRed and the GFP/DsRed ratio of the *MZmir430* mutants for each construct. Data are represented as mean ± SD of biological triplicates. Student *t*-test *P*-values are indicated as *** <0.001 and **** <0.0001.

To understand the effect of microRNA-mediated repression on mRNA structural changes, we analyzed miR-430, which regulates hundreds of maternal mRNAs during the MZT (Fig. 1a, 4a). The miR-430 targets are initially translationally repressed at 4 hpf without significant signs of decay^42^. If ribosomes play a crucial role in shaping mRNA structure *in vivo*, mRNAs targeted by miR-430 should reach an intermediate state at 4 hpf, characterized by an increase in structure due to a depletion of ribosomes. To test this hypothesis, we monitored structural changes in 483 miR-430 targets and a control set of 483 randomly selected mRNAs. We observed a global increase in structure (orange) for miR-430 targets compared to the control set, which is characterized by a decrease in structure (turquoise) between 2 and 4 hpf (Fig. 4d, e). These results support a central role of the ribosome in orchestrating mRNA structure in the cell, suggesting that biological processes regulating translation can have a broad impact on directing mRNA folding.

### 3’-UTRs have a distinct folding landscape

We find that zebrafish transcripts are globally more accessible *in vivo* than *in vitro* (Extended Data Fig. 6a). To investigate whether this global unfolding was uniformly distributed along the transcript, we performed a metagene analysis of each structure, for both *in vivo* and *in vitro* conditions, using a sliding-window approach (Fig. 1b). Our results reveal that regions with decreased structure *in vivo* are mainly found in the 5’-UTR and CDS regions of highly translated transcripts (Fig. 5a, b and Extended Data Fig. 6b), in agreement with the remodeling role of the ribosome. Conversely, 3’-UTRs show increased structure *in vivo*. This increase in RNA structure *in vivo* doesn’t seem to result primarily from RNA binding protein footprints, since no decrease in accessibility was observed over the binding sites of an AU-rich elements binding protein, KHSRP, and surrounding regions recognized by the Ago2/ miR-430 complex (Extended Data Fig. 7). This increase in structure is anti-correlated with translation efficiency, and spreads into the CDS of poorly translated mRNAs, suggesting that the increased structure is specific to ribosome-free regions (Fig. 5b and Extended Data Fig. 6b).

**Figure 6.**
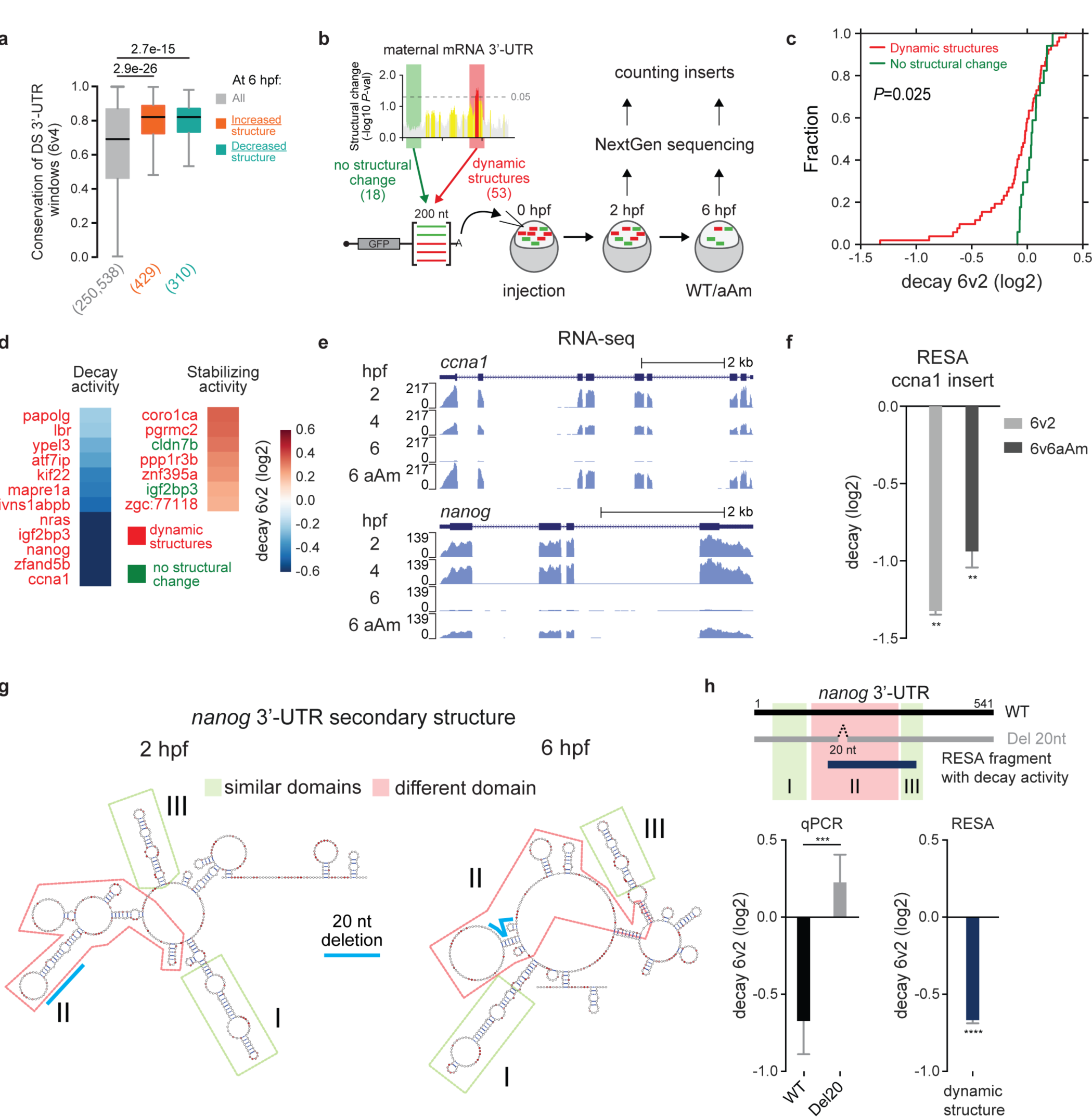
Dynamic 3’-UTR structures are enriched in decay regulatory elements. **a**, Sequence conservation of differentially structured 3’-UTR regions between 4 and 6 hpf (6v4), compared to all 3’-UTR regions analyzed (gray). *P*-values were computed using one-sided Mann-Whitney *U* tests. Box spans first to last quartiles and whiskers represent 1.5× the interquartile range. **b**, Cartoon representation of the RESA experiment used to validate the regulatory activity of 3’-UTR regions that were identified as changing (red) or not changing (green) in their RNA structure during the MZT. **c**, Cumulative distribution of the decay activity measured by the RESA validation experiment, for 3’-UTR regions with dynamic structures (red) or no structural change (green) during the MZT. *P*-values was computed using a one-sided Mann-Whitney *U* test. **d**, RESA-validated 3’-UTR sequences with identified decay or stabilizing elements. **e**, Endogenous mRNA expression levels of *ccna1* and *nanog* at 2, 4 and 6 hpf, as well as for alpha-amanitin-treated (aAm) embryos collected at 6 hpf. **f**, Quantification of the decay activity of *ccna1* measured by RESA, focusing on reporter level changes between 2 and 6 hpf (6v2, light gray), and between 6 hpf untreated and aAm-treated samples (6v6aAm, dark gray). Data are represented as mean ± SD. Student *t*-test *P*-values are indicated as ** <0.01. **g**, Predicted RNA secondary structures of the 542-nt *nanog* 3’-UTR at 2 and 6 hpf. Structural domains that are similar at both developmental stages (I and III) are boxed in green, whereas the one changing (II) is boxed in red. Cyan lines highlight the 20-nt deletion that disrupts a stem region in both structures, which was used for RT-qPCR analysis. **h**, Location of the 20-nt deletion and the 200-nt RESA fragment with decay activity within the *nanog* 3’-UTR (top panel). Decay activity of the *nanog* wild type 3’-UTR (black) and with the 20-nt deletion (gray), quantified by qPCR (6v2), compared to the activity of the 200-nt fragment from the RESA experiment (dark blue). Data are represented as mean ± SD. Student *t*-test *P*-values are indicated as ** <0.01, *** <0.001 and **** <0.0001.

To evaluate the importance of the 3’-UTR RNA folding landscape in the cell, we analyzed the impact of RNA structures on miR-430 activity. Since the accessibility of miRNA binding sites has been shown to affect their activity^9,43,44^, we analyzed how miRNA targeting efficiency correlated to the stability (ΔG) of the predicted structure *in silico, in vitro* and *in vivo* (Fig. 5c, d). We found that the stability of the RNA structure (ΔG) based on *in vivo* probing is the best predictor of miR-430 activity (*rho*=−0.70), outperforming *in silico* (*rho* =-0.39) and *in vitr*o (*rho*=−0.57) structure predictions. Consistent with these results, endogenous miRNA targets with higher free energy *in vivo* were more strongly regulated by miR-430 than those with lower free energy (Fig. 5e, f, i, j and Extended Data Fig. 8). To confirm the regulatory role of those *in vivo* specific RNA conformations, we compared the activity of endogenous targets with different free energy and analyzed the effect that altering their structure had on their regulation. To this end, we quantified the repression of a GFP reporter mRNA in wild type and mutant embryos where the miR-430 locus was deleted (MZ*miR430*). As a control, we co-injected a DsRed reporter mRNA not targeted by miR-430. Modifying the stability of the *in vivo* structure in the target site altered the regulatory strength of each site. For example, destabilizing the *in vivo* structure of the rab33ba target site, which is poorly regulated *in vivo* (Fig. 5f), increased miR-430-mediated regulation (Fig. 5g, h). Conversely, stabilizing the *in vivo* structure of fam171a1, which is strongly regulated by miR-430 (Fig. 5j), abolished miR-430 activity (Fig. 5k, l). Altogether, these results demonstrate that there are increases in 3’-UTR structures *in vivo*, which can regulate gene expression by modulating miRNA activity.

### Dynamic 3’-UTRs enriched in functional elements

During the MZT, maternally-deposited factors (Nanog, Oct4 and SoxB1) mediate activation of the zygotic genome^28^, after which different programs mediate post-transcriptional regulation of maternal mRNAs (Fig. 1a). This process is required for the embryo to transition from the maternal to the zygotic state, and is characterized by changes in the transcriptional and epigenetic landscapes, cellular motility and the cell cycle, with an extension of G1 and G2^27^.

Analysis of the structurally dynamic regions in 3’-UTRs between 4 and 6 hpf revealed an enrichment for conserved sequences (Fig. 6a and Extended Data Fig. 9a), suggesting a potential role in the post-transcriptional regulation of the maternal mRNAs. To test this hypothesis, we performed a parallel reporter assay (RESA)^45^ to measure the regulatory role of 71 different 3’-UTR regions derived from maternal mRNAs subject to decay during the MZT (Fig. 6b). Out of these 71 regions, 53 contained dynamic RNA structures, whereas 18 showed no structural changes during MZT. These sequences, inserted into the 3’-UTR of a reporter gene, were *in vitro* transcribed and injected into 1-cell stage embryos. Then, the abundance of each reporter was measured by high-throughput sequencing at 2 and 6 hpf (Fig. 6b). To identify zygotic-dependent regulation, we compared the levels of each fragment in the presence and the absence of zygotic transcription (i.e. with and without alpha-amanitin). Quantification of mRNA reporters identified decay elements in the 3’-UTR of the *cyclin A1* (*ccna1*) and *nanog* mRNAs and an enrichment of destabilizing elements in regions that change in structure during development compared to regions that do not change (*P*=0.025) (Fig. 6c, d, and >Extended Data Fig. 9b-e). These results suggest that regions with dynamic structures are enriched in functional regulatory elements.

One such dynamic regulatory region is located in the *ccna1* mRNA. Cyclin A1 controls the cell cycle and is actively degraded during the MZT. Time-course analyses of mRNA expression during the MZT using RNA-seq reveal that *ccna1* mRNA decays in wild type embryos but not in embryos where zygotic transcription is blocked with alpha-amanitin (Fig. 6e and Extended Data Fig. 9f). The structurally dynamic region of *ccna1* tested in RESA also possesses a decay activity that relies on the activation of the zygotic program (Fig. 6f). This is consistent with the regulation of *ccna1 in vivo*, and in the change in cell cycle length after zygotic genome activation. Another dynamic region with regulatory activity resides in the *nanog* mRNA, a transcription factor required for genome activation and miR-430 expression, and which is degraded after zygotic genome activation (Fig. 6e and Extended Data Fig. 9f). We predicted the secondary structure of the *nanog* 3’-UTR at 2 and 6 hpf, using *in vivo* DMS-seq accessibilities as constraints (Fig. 6g). We identified two domains with constant structure (I and III), flanking a region with differential structure (II). This domain (II) overlaps with the dynamic region that had regulatory activity *in vivo* (Fig. 6d, h). To test the regulatory activity of this region in the context of the 3’-UTR, we compared the regulation provided by the wild type *nanog* 3’-UTR to a deletion mutant reporter that disrupted the stem region (II) with a 20-nt deletion (Fig. 6g, h). The full-length *nanog* 3’-UTR had a decay activity similar to that of the differentially structured region tested in RESA (Fig. 6h). However, a deletion of 20 nucleotides in domain II stabilized the reporter mRNA, suggesting that this region is required to confer regulation *in vivo* (Fig. 6g, h). Altogether, these results demonstrate that tracking mRNA structure dynamics throughout biological transitions can reveal functional elements in 3’-UTRs.

## Discussion

Here we examine how RNA structure dynamics change during the MZT, a universal regulatory transition in animal embryogenesis. Our study reveals that 3’-UTRs and untranslated mRNAs are more structured in the embryo than *in vitro.* Dynamic 3’-UTR structures *in vivo* correspond to novel regulatory regions controlling maternal mRNA decay. Lastly, we show that the ribosome is one of the principal factors unwinding the mRNA, and while structure does not emerge as a global regulator of translation, translation is a major driver shaping the mRNA structure landscape during early embryogenesis.

Our demonstration that untranslated mRNAs and 3’-UTRs are more structured *in vivo* than *in vitro* (Fig. 5a, b and Extended Data Fig. 6b) runs contrary to work showing that mRNAs are mostly unfolded in the cell^13,15^. Previous *in vitro* and *in vivo* analyses of RNA structures have been carried out in yeast, where the average 3’-UTR length is much shorter than that of vertebrate mRNAs^13^. In the absence of constitutive unwinding by the ribosome, both molecular crowding and RNA binding proteins are likely to favor higher-order RNA structures^46,47^. Remodeling of mRNA structure can also affect regulatory structural elements modulating 3’-end processing and mRNA stability^48^. Our data support a view where cellular factors explicitly affect 3’-UTR structures, modulating the regulatory activity of miRNAs and gene expression during vertebrate development.

Our analysis of dynamic mRNA structures identifies regulatory regions during the MZT in the *cyclin A1* and *nanog* mRNAs, two maternal factors that are required to regulate the cell cycle and zygotic transcription, and that are rapidly degraded during the MZT. We find that the characterization of structurally dynamic regions during cellular or developmental transitions represents a means of identifying potential regulatory elements *in vivo* (Fig. 6b-d and Extended Data Fig. 9b, c).

Global RNA unwinding in yeast is an active process^13^. The cellular factors responsible for this structural remodeling and the specific regions that are remodeled remain unknown. While the ribosome has been shown to have an helicase activity *in vitro*^16^, previous experiments in exponentially growing yeast do not reveal a relationship between translation and RNA unfolding^13^. Here, we have followed the dynamics of mRNA and show that the ribosome is a major engine for mRNA structure remodeling, promoting alternative RNA conformations for thousands of endogenous transcripts in vertebrate cells. This effect is also observed in 5’-UTRs containing translated uORFs, indicating that the ribosome helicase activity has the potential to regulate elements that rely on specific RNA conformations^49,50^. Several mechanisms regulating mRNA translation, such as changes in poly(A) tail length and microRNA-mediated repression, also result in the remodeling of mRNA structure during embryogenesis.

RNA structure has generally been thought to have a major effect on mRNA translation rates^19^, based on early experiments in bacteria and eukaryotes, where strong hairpin structures in mRNAs reduce translation^20–23^. Here, we have demonstrated precisely the opposite effect, i.e. translation modulates mRNA structure, and mRNA structure does not have a global effect on translation during embryogenesis (Fig. 2 and Extended Data Fig. 2). While individual examples in eukaryotes show that stable 5’-UTR structures can block translation^20,22^, at the level of the transcriptome, complex eukaryotic cells have efficient helicase activities that greatly diminish the global effect of 5’-UTR structure on translation. We propose that eukaryotic systems have evolved powerful helicases that, together with sophisticated mechanisms to regulate translation, such as uORFs, miRNAs, RNA-binding proteins, and longer 3’-UTRs, weaken the transcriptome-wide effects that 5’-UTR and CDS structures might have on translation. Our results provide a conceptual shift in our understanding of the relationship between mRNA structure and translation, and suggest that on a global level, translation guides structure rather than structure guiding translation.

**Supplementary Information** is available in the online version of the paper.

## Acknowledgements

We thank K. Bilguvar, S. Mane and C. Castaldi for sequencing support. We thank A. Bazzini and members of the Giraldez and the Kellis lab for intellectual and technical support. This research was supported by the Fonds de Recherche du Québec - Santé (postdoctoral fellowship to JDB), the Human Frontier Science Program (LT000307/2013-L to EMN), the Australian Research Council (DE170100506 to EMN) and the National Institute of Health (grants R01 HD074078, GM103789, GM102251, GM101108 and GM081602), Pew Scholars Program in the Biomedical Sciences, March of Dimes 1-FY12-230, the Yale Scholars Program and Whitman fellowship funds provided by E. E. Just, Lucy B. Lemann, Evelyn and Melvin Spiegel, The H. Keffer Hartline and Edward F. MacNichol, Jr. of the Marine Biological Laboratory in Woods Hole, MA to AJG.

## Author Contributions

JDB and AJG conceived the project. JDB performed the experiments. JDB, EMN and CEV performed data processing. VY identified regulatory element in the nanog 3’-UTR and built the reporter constructs. CT performed the KHSRP iCLIP experiment. JDB and EMN performed data analysis and, together with AJG, interpreted the results. AJG supervised the project, with the contribution of MK. JDB, EMN and AJG wrote the manuscript with input from the other authors.

## Author Information

Correspondence and requests for materials should be addressed to JDB and AJG.

---

**Extended Data Figure 1.**
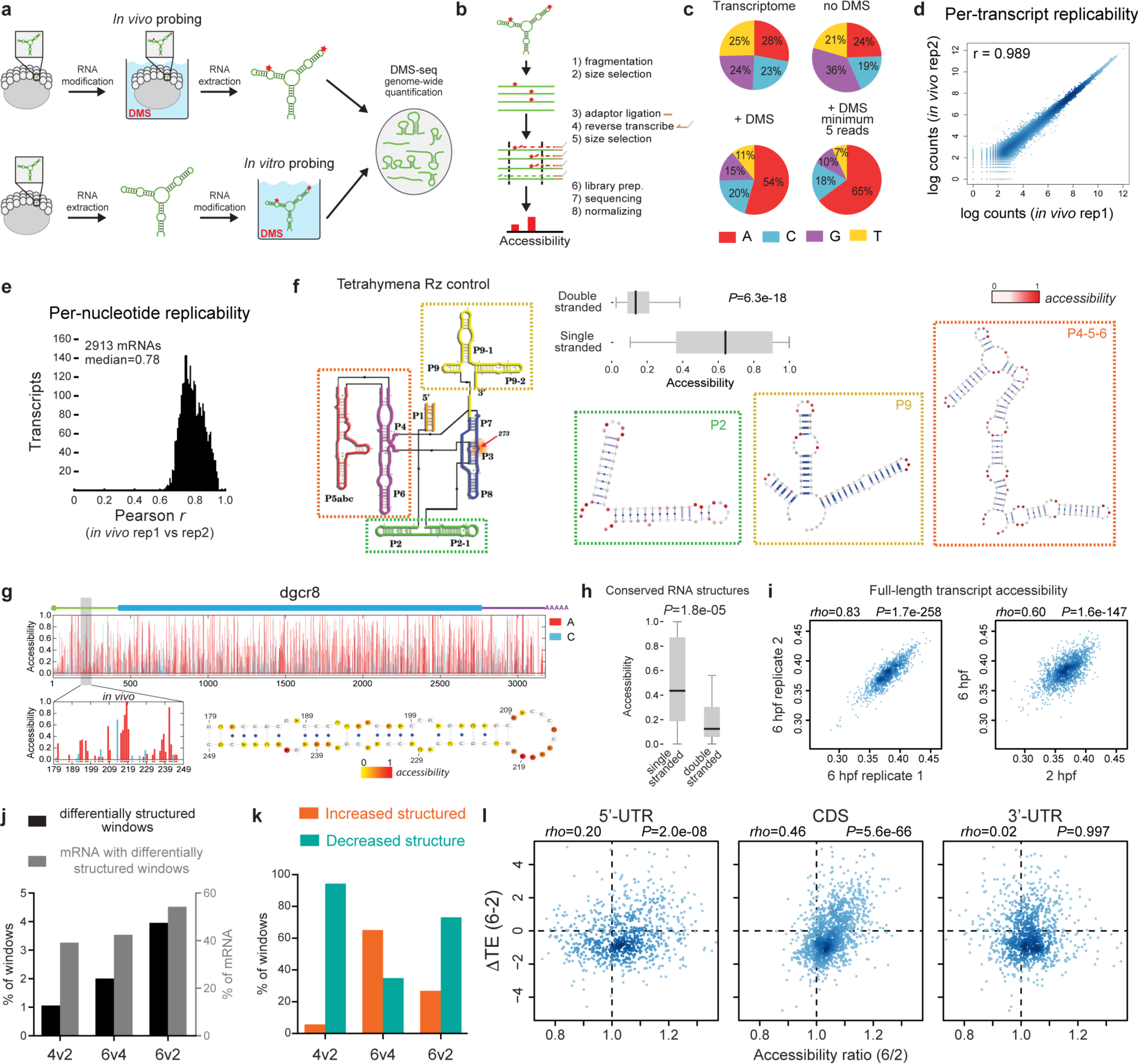
DMS-seq controls and mRNA structure changes during the MZT. To determine how mRNA structure changes during cellular transitions *in vivo*, we analyzed the structure of the zebrafish transcriptome during the MZT using DMS-seq (at 2, 4 and 6 hours post-fertilization, hpf). DMS methylates accessible N1 of adenine and N3 of cytidine bases in single-stranded regions, at the end of a stem, and at base pairs flanking G-U wobble interactions^67^, causing reverse transcriptase drop-off during reverse transcription. The truncated RNAs are then captured, analyzed by high-throughput sequencing and normalized per-transcript, providing per-nucleotide accessibility values between 0 and 1 (see Methods). Three lines of evidence indicate that we can effectively probe RNA structure during embryogenesis. First, we observed a robust and reproducible enrichment of reads mapping to A and C bases (74-83%) compared to DMS-untreated samples (43%) (**c**), as well as reproducible counts between biological replicates (*r* of 0.989) (**d, e**). Second, the DMS-seq accessibility profile of the *Tetrahymena* ribozyme, used as an exogenous control, was in close agreement (*P*=6.3e-18) with the previously reported structure^68^ (**f**). Third, analysis of conserved secondary structures, such as those found in *dgcr8*^69^, *selt1a* and *selt2* mRNAs showed that paired A and C bases exhibited lower accessibilities than single stranded regions (*P*=1.8e-05) (**g, h**). These results indicate that our DMS-seq analysis provides a robust transcriptome-wide map of RNA structure dynamics in a vertebrate embryo. **a**, Schematic view of *in vivo* and *in vitro* genome-wide DMS probing. Red stars represent DMS-modified nucleotides. **b**, Cartoon shows the DMS-seq protocol. Dashed lines highlight the second size selection capturing reverse transcription products that prematurely stop due to DMS modifications. **c**, Proportion of DMS-seq reads from DMS-modified nucleotides *in vivo* at 2 hpf (+DMS), compared to untreated (no DMS) and the composition of the whole transcriptome. *In vivo* DMS treatment markedly enriches A/C modified sites, and is increased when nucleotides with minimum coverage of 5 RT-stops/nt are considered (see Methods). **d**, **e**, DMS-seq data is reproducible across biological replicates at a per-transcript (**d**) and per-nucleotide (**e**) resolution *in vivo* at 2 hpf. **f**, Validated secondary structure of the *Tetrahymena* ribozyme (Rz) (adapted from Walter *et al.*^70^). In the zoomed panels (corresponding to regions P2, P9 and P4-5-6), DMS-seq accessibilities have been overlaid (from white to red) onto the known secondary structures. In the upper middle part of the figure, boxplot represents consistency between accessibilities and Rz pairing status. *P*-value computed using a one-sided Mann-Whitney *U* test. Box spans first to last quartiles and whiskers represent 1.5× the interquartile range. **g**, *In vivo* accessibility profile of the dgcr8 mRNA. Red and turquoise bars correspond to A and C bases, respectively. Accessibilities corresponding to annotated conserved secondary structures (gray rectangles) are highlighted. The detailed secondary structure is also shown, where *in vivo* accessibilities have been overlaid for A and C bases (from yellow to red), while Gs and Us are depicted in white. **h**, Box plot comparing the distribution of *in vivo* accessibilities and the pairing status of A and C bases found in three conserved secondary structures used as controls (selt1a, selt2 and dgcr8). *P*-value computed using a one-sided Mann-Whitney *U* test. Box spans first to last quartiles and whiskers represent 1.5× the interquartile range. **i**, Comparisons of per-transcript DMS-seq accessibilities between replicates (left) and across developmental stages (right). **j**, Proportion of differentially structured windows (see Methods) across the 3 developmental stages, expressed as the percentage of differentially structured windows-relative to the total windows analysed-(black), and the percentage of mRNAs that contain these windows –relative to the total transcripts present in both stages-(gray). **k**, Proportion of differentially structured windows that increase (orange) or decrease (turquoise) in RNA structure, for each pairwise comparison, based on the directionality of their change in Gini index. The direction of change is defined with respect to the latest developmental stage, for each pairwise comparison. **l**, Correlation between the per-transcript changes in translation efficiency and accessibility for each transcript region (5’-UTR, CDS and 3’-UTR) between 2 and 6 hpf.

**Extended Data Figure 2.**
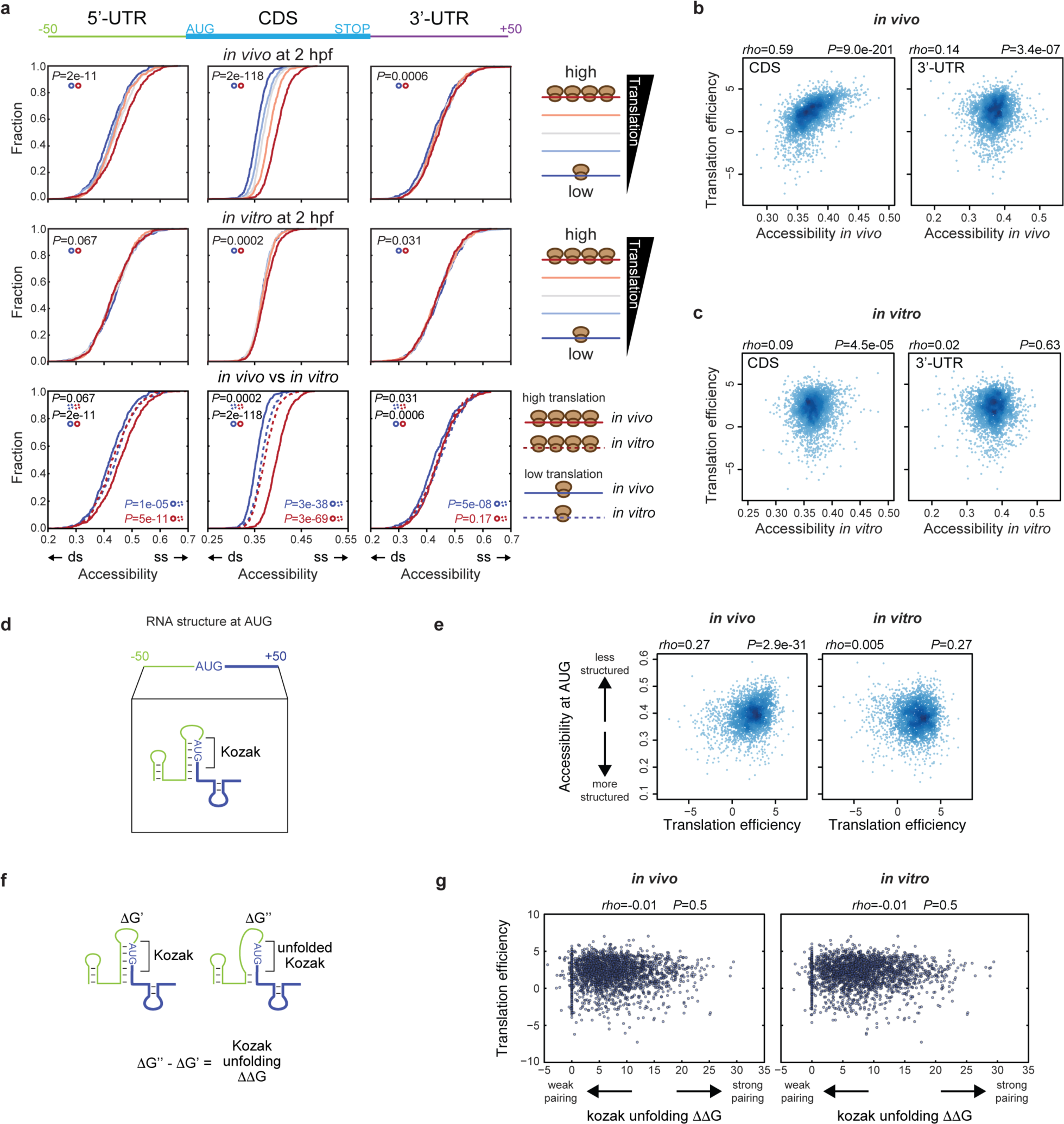
Ribosomes unwind RNA structures in 5’-UTR and CDS, including the region surrounding the AUG. **a**, Cumulative distributions of global accessibilities of the 50 nucleotides upstream of the AUG, CDS or 50 nucleotides downstream of the STOP codon are shown for *in vivo* (top), *in vitro* (middle) and *in vivo* vs *in vitro* (bottom) samples at 2 hpf. mRNAs have been binned in quintiles according to their translation efficiency (TE) and colored from blue (lowest TE) to red (highest TE). *P*-values have been computed using a two-sided Wilcoxon signed-rank test and a one-sided Mann-Whitney *U* test, for comparisons within and across translational statuses, respectively. **b**, **c**, Correlation between translation efficiency and the accessibility of CDS (left) and 3’-UTR regions (right), for both *in vivo* (**b**) and *in vitro* (**c**) conditions. **d**, Cartoon shows the region surrounding the AUG initiation codon used to predict RNA secondary structure and highlights the Kozak region (5’-NNNNAUGNNN-3’). **e**, Correlation between translation efficiency and the accessibility at the AUG initiation codon for both *in vivo* (left) and *in vitro* (right) conditions. **f**, The RNA secondary structure and the Gibbs free energy (ΔG’) were computed for each AUG initiation codon. The energy required to unfold the Kozak region (ΔΔG) was calculated by subtracting the ΔG’ from the energy of the structure where the Kozak sequence is forced to be single-stranded (ΔG’’). **g**, Correlation between translation efficiency and the energy needed to unfold and make accessible the Kozak sequence (ΔΔG) for both *in vivo* (left) and *in vitro* (right) conditions. No correlation is observed in both cases, suggesting that the structure of the Kozak sequence is not a major factor driving translation in the early embryo.

**Extended Data Figure 3.**
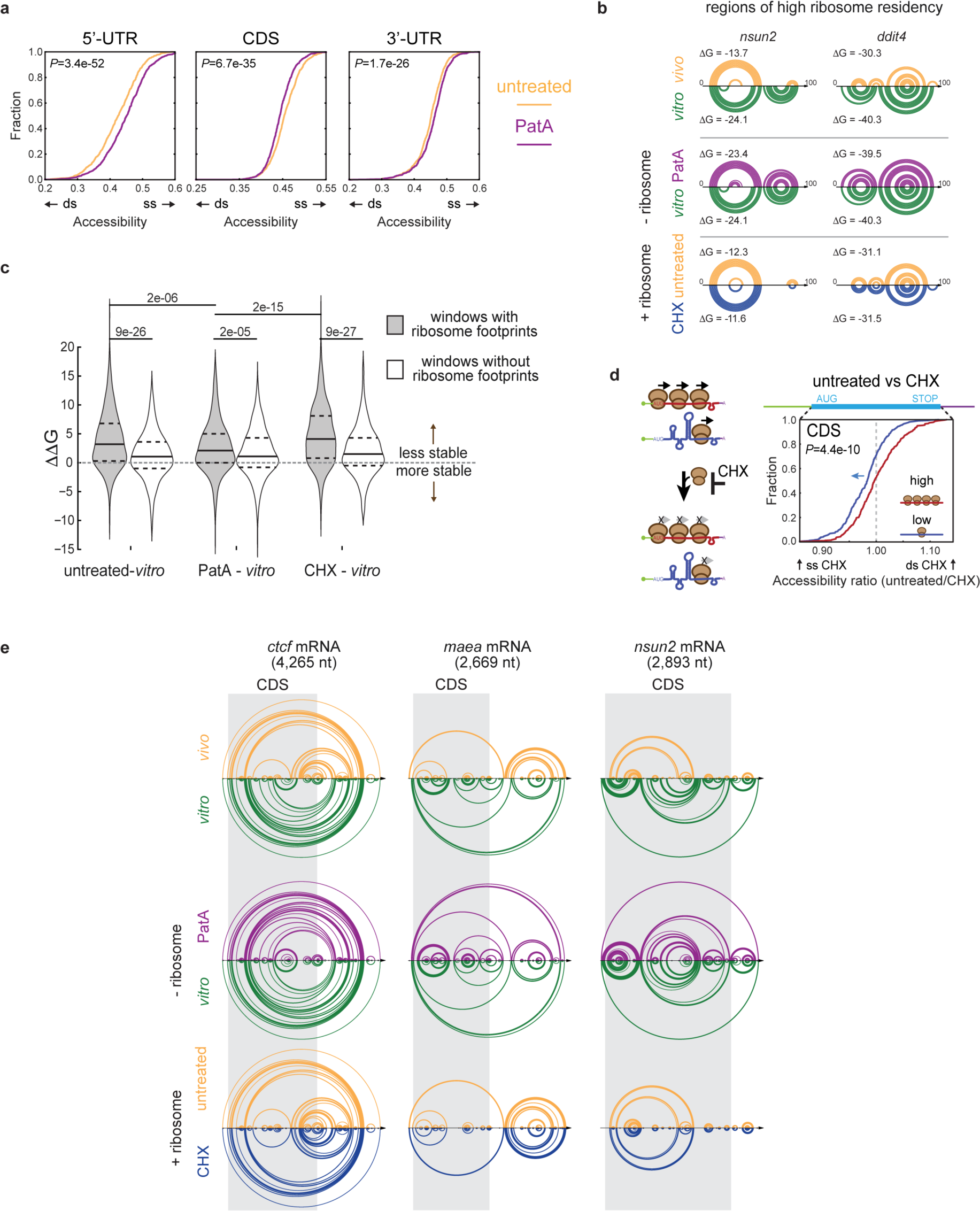
Ribosomes promote alternative RNA conformations with reduced stability in the cell. **a**, Cumulative distributions of 5’-UTR, CDS and 3’-UTR global accessibilities in PatA-treated and untreated samples. *P*-values have been computed using a two-sided Wilcoxon signed-rank test. **b**, Arc plots of DMS-seq guided RNA secondary structure predictions of high ribosome occupancy regions found in the *nsun2* and *ddit4* genes, for *in vivo* (yellow) *in vitro* (green), untreated (yellow), PatA-treated (purple) or CHX-treated (blue) samples. Each arc represents a base pair interaction. **c**, Pairwise comparisons of Gibbs free energy differences (ΔΔG; untreated-*in vitro*, PatA-*in vitro* and CHX-*in vitro*) of windows with high ribosome footprint densities (gray) and without ribosome footprints (white). Positive ΔΔG indicates a less stable structure in the first condition (untreated, PatA or CHX), compared to *in vitro*, whereas negative ΔΔG indicates a more stable structure in the first condition, compared to *in vitro*. Note the decrease in stability for structures formed by ribosome-rich regions (gray) in the untreated and CHX-treated samples, but not in the ribosome-free PatA-treated sample or in regions without ribosome (white). Violin plot features a kernel density estimation and lines represent the quartiles of the distribution. *P*-values have been computed using a two-sided Wilcoxon signed-rank test and a one-sided Mann-Whitney *U* test, for comparisons across probing conditions for a same group and between groups within a specific condition, respectively. **d**, Cartoon representation depicting the impact of CHX, along with its corresponding cumulative distribution of CDS accessibility ratios (untreated/CHX) for highly and lowly translated mRNAs at 2 hpf. Note the minor increase in accessibility in lowly translated mRNAs following CHX treatment (blue arrow). P-value was determined using a one-sided Mann-Whitney *U* test. **e**, Arc plots of DMS-seq guided RNA secondary structure predictions of the full-length *ctcf, maea* and *nsun2* mRNAs using SeqFold^40^. The CDS region has been shadowed in gray.

**Extended Data Figure 4.**
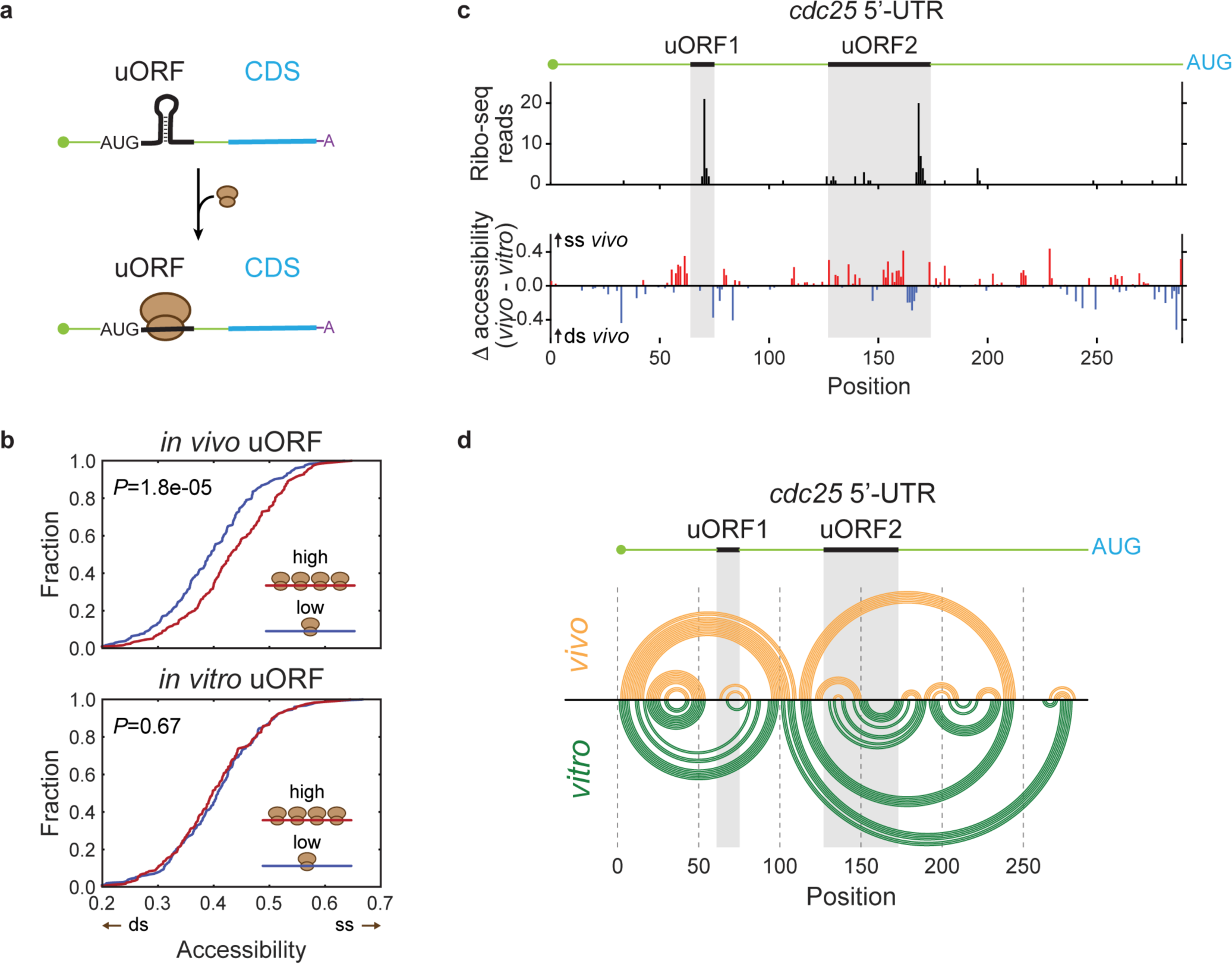
uORF translation remodels 5’-UTR structure. **a**, Schematic view of the 5’-UTR RNA structure remodeling that occurs upon translation of upstream open reading frames (uORF). **b**, Cumulative distributions of *in vivo* (top) and *in vitro* (bottom) uORF accessibilities for highly (red) and lowly (blue) translated uORFs. *P*-values were determined using one-sided Mann-Whitney *U* tests. **c**, Distributions of Ribo-seq reads (top panel) and accessibility differences (*in vivo* – *in vitro*) (bottom panel) for the *cdc25* 5’-UTR, which contains two translated uORFs (highlighted in gray). Increased accessibility *in vivo* is shown in red, whereas decreased accessibility *in vivo* is shown in blue. **d**, Arc plot depicting the predicted *cdc25* 5’-UTR secondary structure, guided by either *in vivo* (yellow) or *in vitro* (green) DMS-seq accessibilities. Each arc represents a base pair interaction.

**Extended Data Figure 5.**
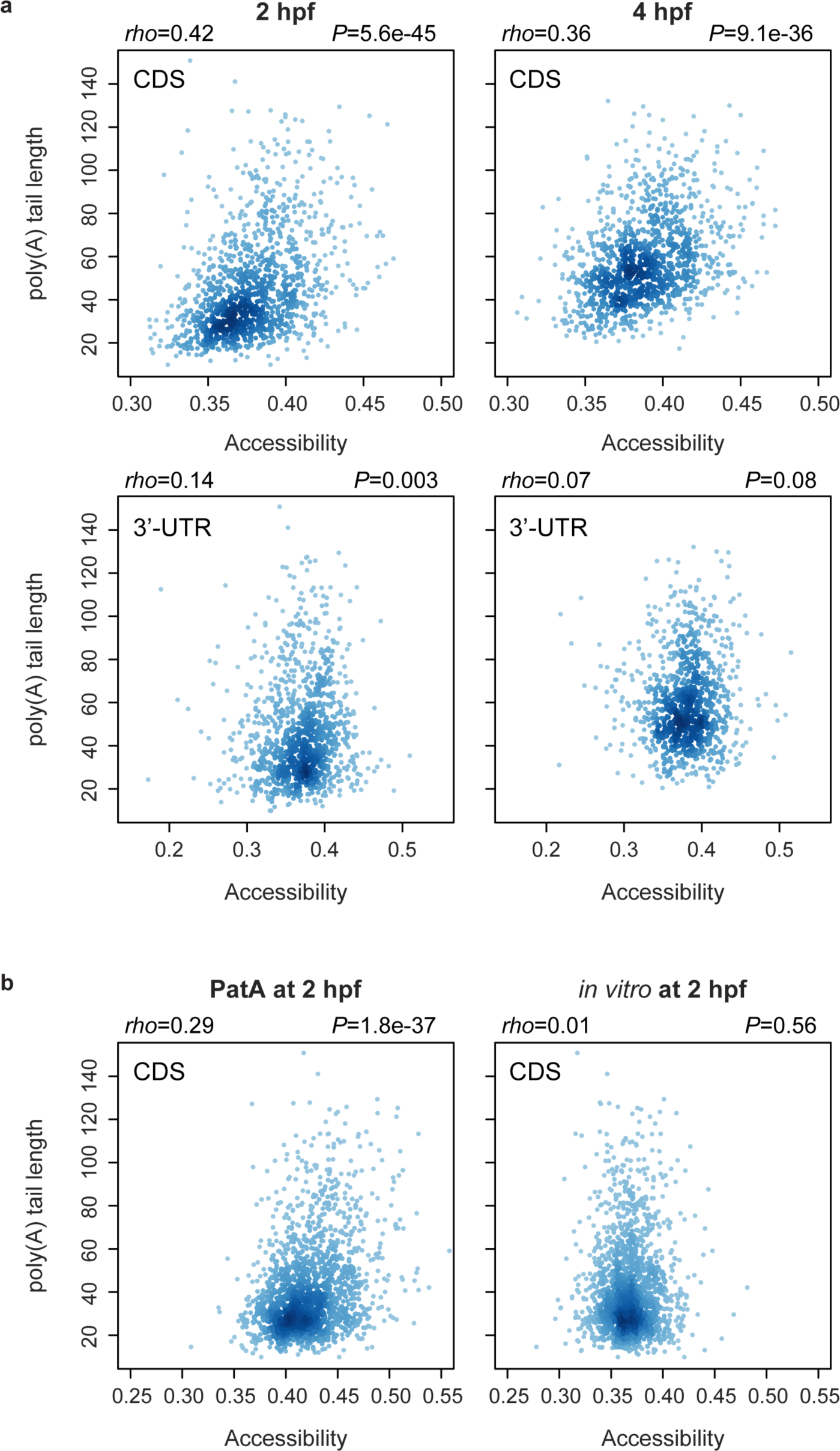
Poly(A) tail length-dependent mRNA structure remodeling depends on translation. **a**, Comparison of per-region accessibilities (CDS and 3’-UTR) and per-transcript poly(A) tail length, at 2 hpf (left panels) and 4 hpf (right panels). **b**, Comparison of CDS accessibilities and per-transcript poly(A) tail lengths in PatA-treated (left) and *in vitro* refolded samples (right) from 2 hpf embryos.

**Extended Data Figure 6.**
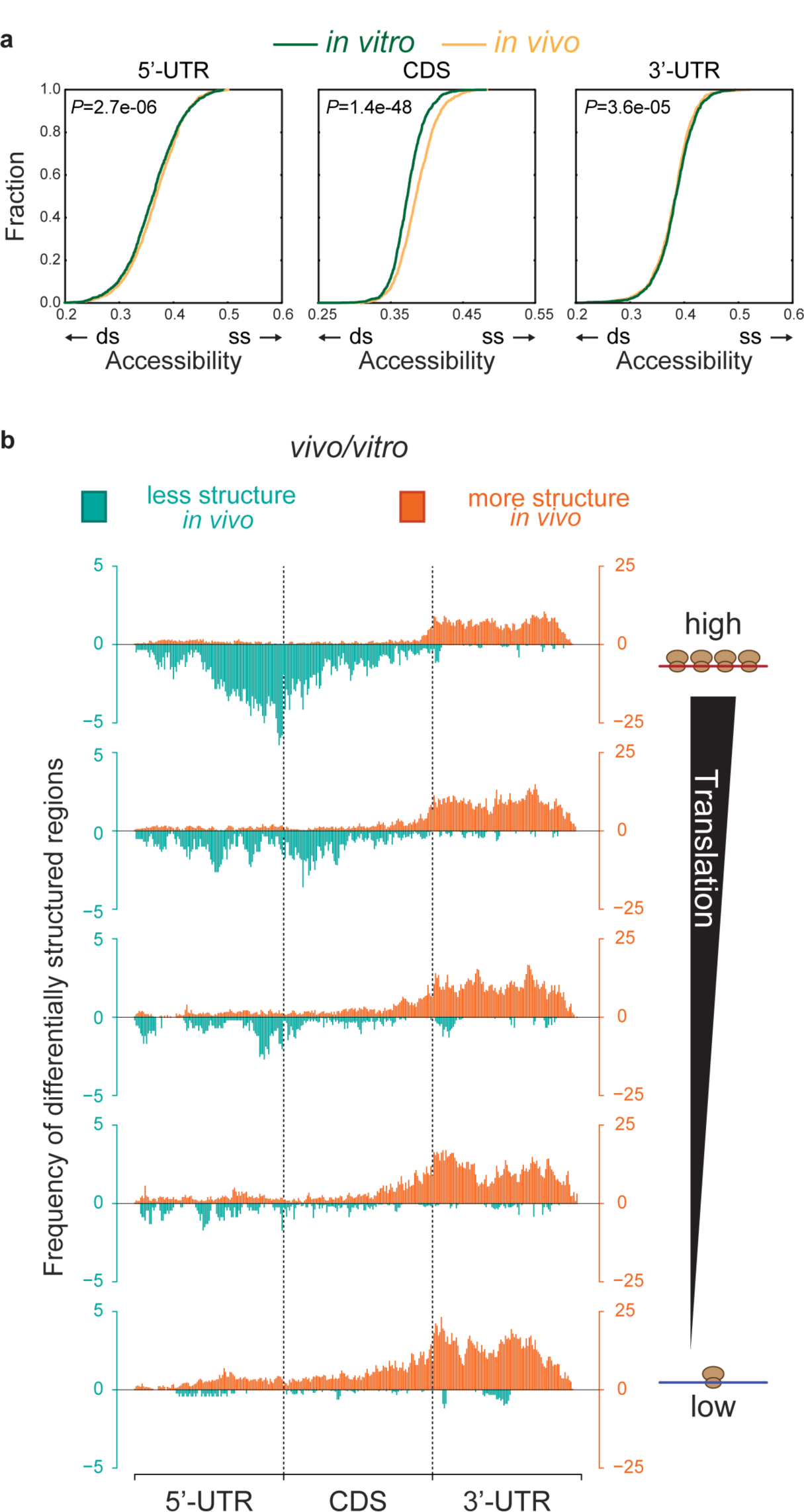
Differences between *in vivo* and *in vitro* RNA structures are not uniformly distributed along transcripts. **a**, Cumulative distributions of global 5’-UTR, CDS and 3’-UTR accessibilities *in vivo* and *in vitro*. Only transcripts with a minimum of 85% coverage of As and Cs and with at least 10 reads in average per As and Cs for both *in vivo* and *in vitro* DMS-seq experiments are shown. *P*-values were computed using two-sided Wilcoxon signed-rank tests. **b**, Distribution of differentially structured windows along the transcripts, comparing *in vivo* and *in vitro* conditions. Windows with increased structure *in vivo* are depicted in orange, whereas those with decreased structure *in vivo* are shown in turquoise. Each transcript region (5’-UTR, CDS, 3’-UTR) has been normalized by its length, as well as by total number of windows analyzed in each region. Transcripts have been binned into quintiles based on their translation efficiency, which has been determined using ribosome profiling data.

**Extended Data Figure 7.**
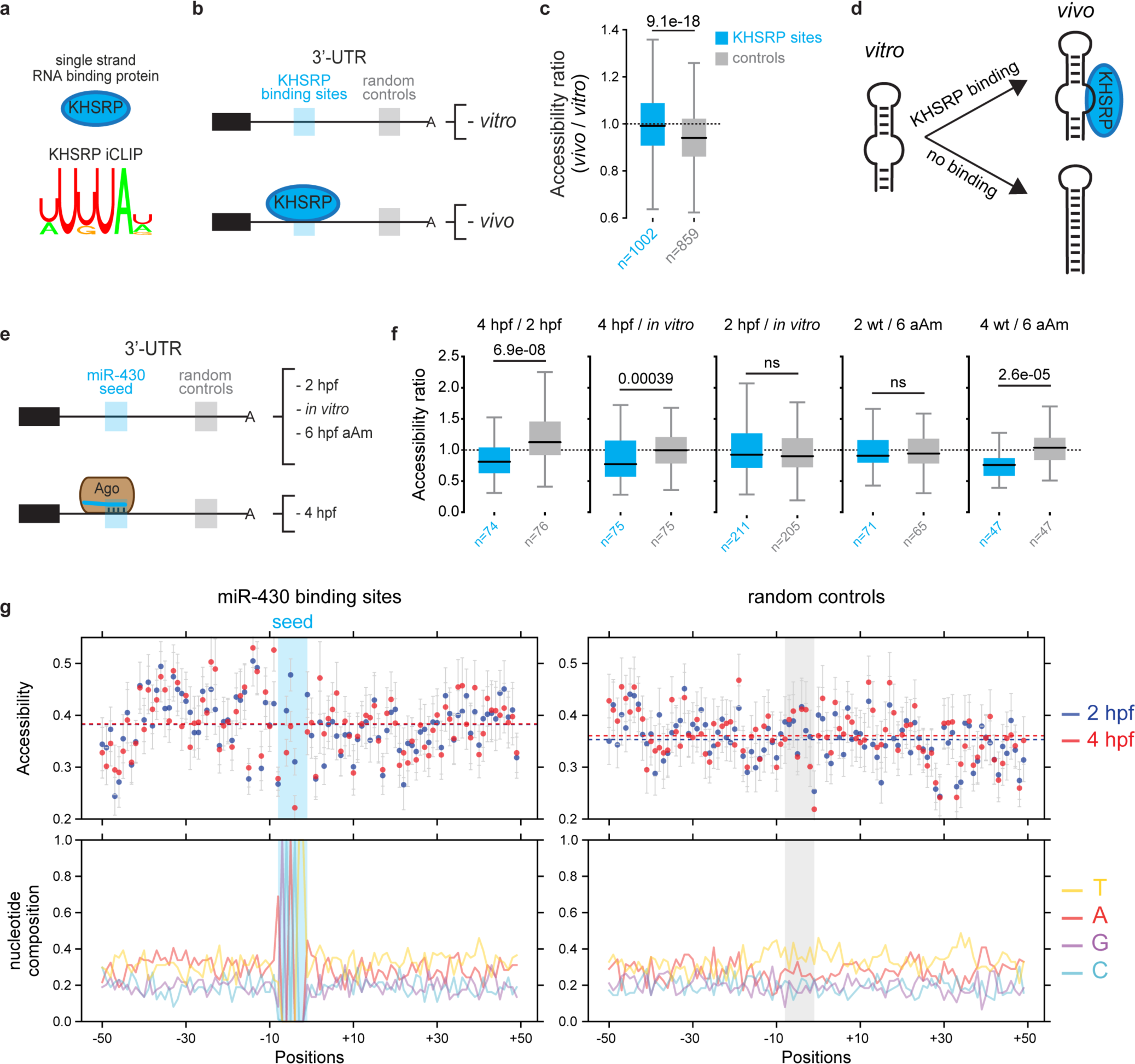
DMS-seq signal provides information on the RNA conformation favored upon the binding of two different *trans*-factors. While a high DMS-seq signal implies a single-stranded conformation, a low DMS-seq signal signifies either an RNA-RNA or RNA-protein interactions involving the Watson-Crick face of the base^71^. To test whether DMS probing is mainly analyzing RNA-RNA pairing *versus* protein footprinting *in vivo*, we analyzed the structure over two types of regions, one bound by the RNA binding protein KHSRP identified by iCLIP and another occupied by the Ago2/miR-430 complex. We find that KHSRP-bound 3’-UTR regions are more accessible *in vivo*, compared to control regions from the same transcripts, and comparable to *in vitro*, suggesting that KHSRP binding helps to maintain a single-stranded RNA conformation in the cell. In addition, decreased accessibility was observed at miR-430 seed regions when the Ago2/miR-430 complex is present, but not in the flanking sequences nor in absence of miR-430 expression, therefore measuring seed-miRNA interactions. Analyses from two different types of *trans*-factors (KHSRP and Ago2/miR-430 complex) suggest that our approach measures the RNA structure promoted by those factors rather than their footprints limiting DMS’s access to RNA. **a**, Sequence logo of KHSRP identified using iCLIP. **b**, Cartoon depicting the binding status of KHSRP to its target *in vitro* and *in vivo*. **c**, Comparison of the accessibility ratio (*in vivo* / *in vitro*) of KHSRP-bound regions (cyan) and matching unbound controls (gray), suggesting that KHSRP binding helps to maintain a single-stranded RNA conformation in the cell. KHSRP binding sites were determined using iCLIP experiments (see Methods) and limited to those found within 3’-UTRs. Unbound control regions originate from 3’-UTR targeted by KHSRP, overlap with less than 25% of the binding site and possess the same length than the binding site. *P*-value computed using one-sided Mann-Whitney *U* test. Box spans first to last quartiles and whiskers represent 1.5× the interquartile range. **d**, Schematic representation of the impact of KHSRP binding on RNA structure *in vivo* keeping bound-RNA in a single-stranded conformation. **e**, Cartoon depicting the binding status of the Ago2/miR-430 complex to its target in various conditions. **f**, Comparison of accessibility ratios (across conditions and time points) of miR-430 seeds (cyan) and matching controls (gray). miR-430 seeds correspond to any 8- and 7-mers miR-430 binding sites found in the 3’-UTR of miR-430 targets (see Methods). Control regions of 8-nt were randomly chosen within the miR-430 seed containing 3’-UTRs (see Methods). MiR-430 seed and control accessibilities were calculated by averaging the accessibility of A and C bases found in each 8-nt sequence. *P*-values computed using one-sided Mann-Whitney *U* tests. Box spans first to last quartiles and whiskers represent 1.5× the interquartile range **g**, Meta-analysis of average accessibilities at 2 and 4 hpf (top panels) and nucleotide content (bottom panels) at each position of miR-430 seed (n=74, left panels) and control (n=75, right panels) regions. Each region corresponds to a 100-nt window centered on a 8-nt miR-430 seed or control sequences highlighted in cyan and gray, respectively. Dotted lines correspond to the average accessibility of the metaplot for each developmental stage. Error bars denote s.e.m.

**Extended Data Figure 8.**
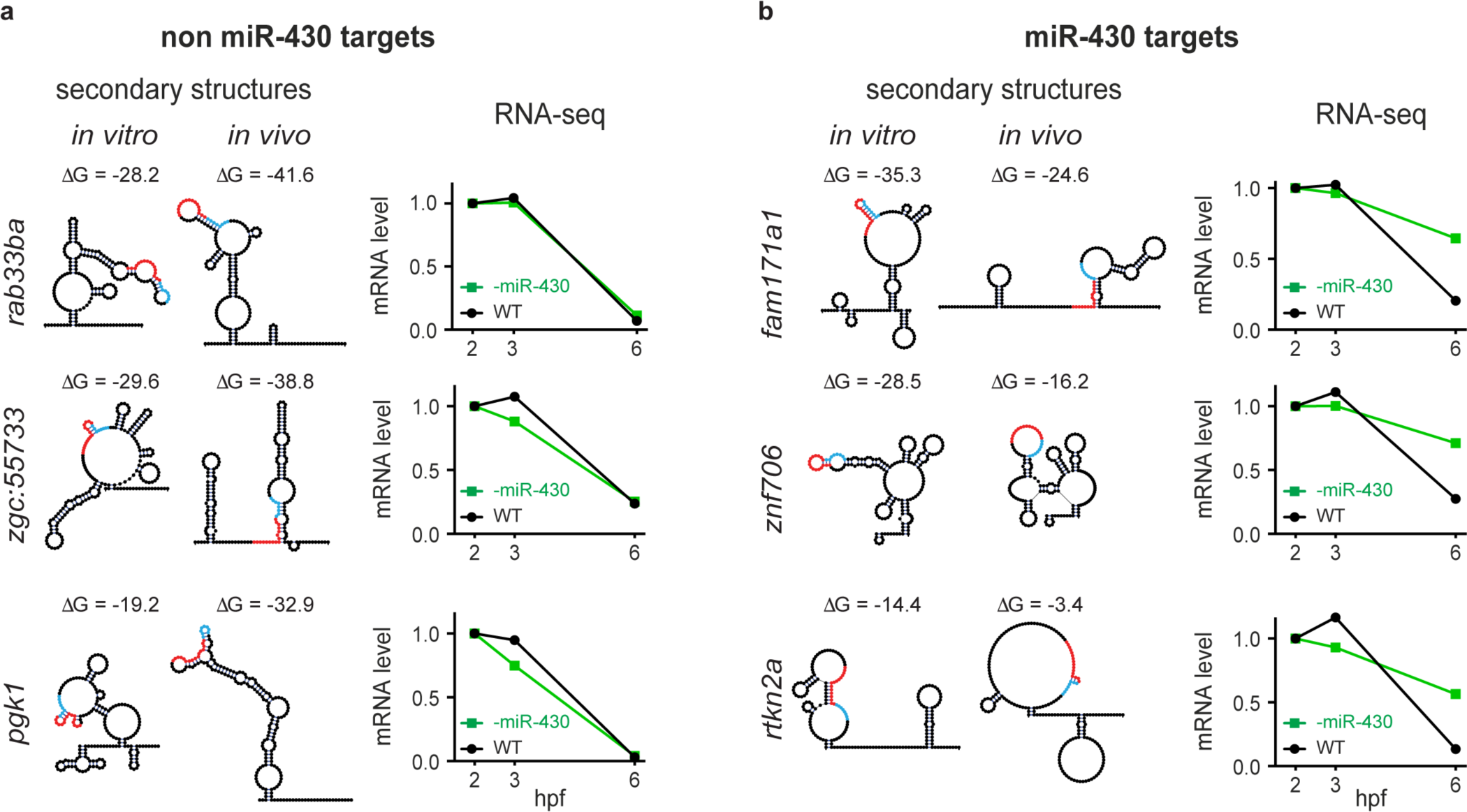
*In vivo* specific 3’-UTR structures impact miRNA activity and gene expression. **a**, **b**, *In vivo* and *in vitro* predicted secondary structures and stabilities (ΔG) of the 200-nt region centered on the miR430 target site found in non miR-430 targets (*rab33ba, zgc:55733* and *pgk1*; **a**) and miR-430 targets (*fam171a1, znf706* and *rtkn2a*; **b**). Regions complementary to the miR-430 seed and its complementary binding region within the 3’-UTR are shown in cyan and red, respectively. Changes in mRNA abundance during the MZT of the endogenous transcripts in wild type conditions (black) or in conditions where miR-430 activity is inhibited by a tinyLNA complementary to the miR-430 seed (green) are shown for each gene. MiR-430 targets are characterized by a gain in stability at 6 hpf when miR-430 activity is reduced (-miR-430) (**b**).

**Extended Data Figure 9.**
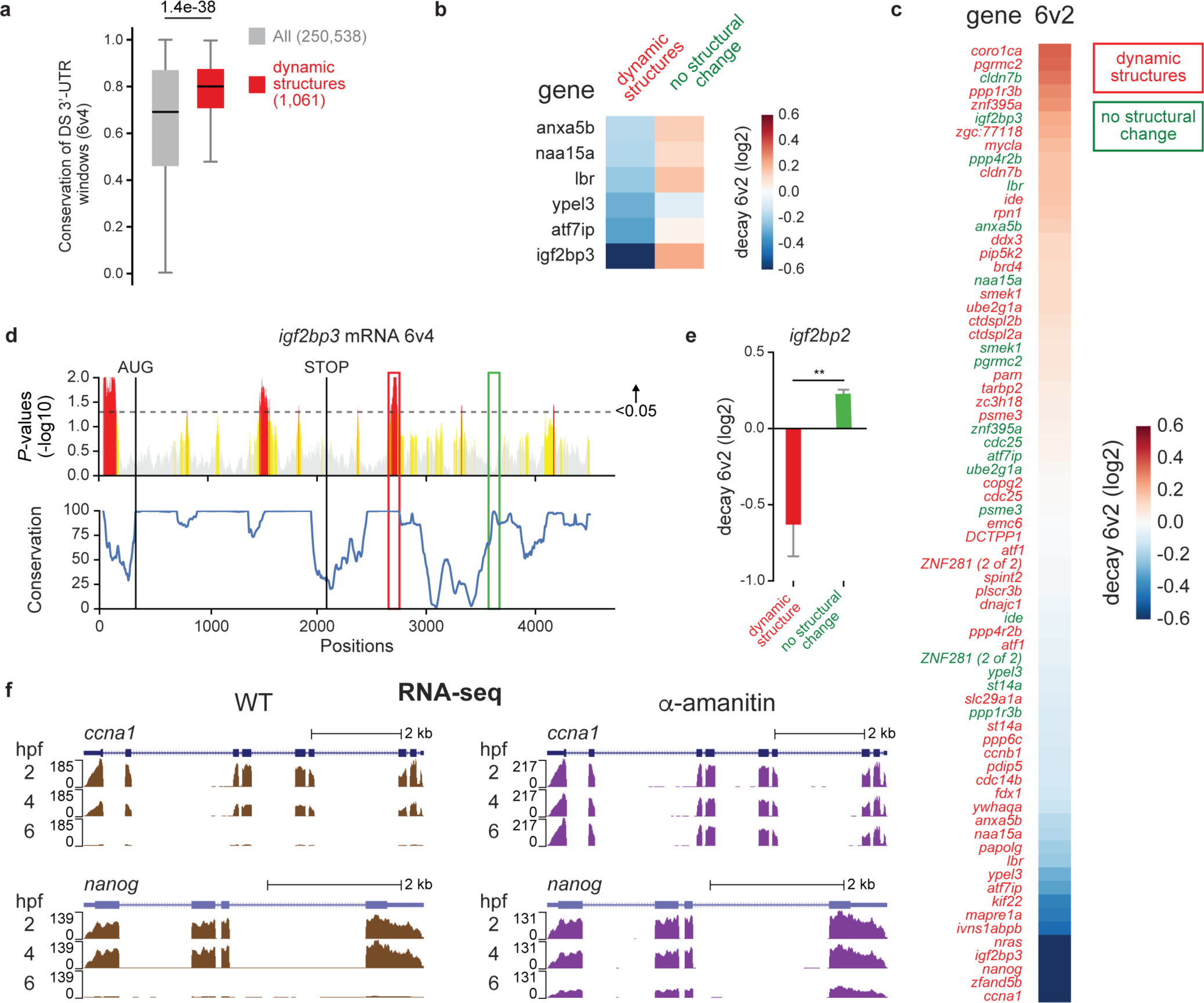
Dynamic 3’-UTR structures reveal decay elements during the MZT. **a**, Enrichment of conserved sequences in 3’-UTR regions with a dynamic structure between 4 and 6 hpf (6v4), compared to all 3’-UTR regions analyzed. Dynamic structures correspond to those with a KS test *P*-value <0.05. Box spans first to last quartiles and whiskers represent 1.5× the interquartile range. *P*-value computed using a one-sided Mann-Whitney *U* test. **b**, Decay activity of regions with a dynamic structure (left column, red) and a static structure (right column, green) originated from the same 3’-UTR, calculated from the RESA experiment. **c**, Decay (blue) and stabilizing (dark orange) activities of all the 200-nt 3’-UTR regions with sufficient read coverage from the RESA experiment (see Methods). Genes labeled in red correspond to those that have regions with a dynamic structure during the MZT, whereas genes labeled in green correspond to those with no structural change. Note the enrichment of regions with a dynamic structure among the decay elements (blue). **d**, Per-transcript KS test *P*-value profile (top) and conservation (bottom) for the *igf2bp3* mRNA, comparing 4 and 6 hpf. The profiles have been computed by analyzing 100-nt sliding windows throughout the transcript. Examples of conserved 3’-UTR regions with a dynamic structure (red) and without structural change (green) are highlighted. *P*-values between 1 and 0.2 are shown in gray, between 0.2 and 0.05 are shown in yellow and <0.05 are shown in red. The horizontal dotted gray line indicates *P*-value equal to 0.05. The vertical black lines delineate the AUG and STOP codons of the CDS. **e**, Quantification of the decay activity for the dynamic (red) and non-changing (green) structures found in the *igf2bp3* 3’-UTR, as calculated by RESA. Data are represented as the mean ± SD. Student *t*-test *P*-values are indicated as ** <0.01. **f**, Genome tracks of RNA-seq experiments representing mRNA levels of the *ccna1* (top) and *nanog* (bottom) transcripts at 2, 4 and 6 hpf in wild type (left) and alpha-amanitin (right) conditions. Alpha-amanitin inhibits zygotic genome activation, highlighting the implication of zygotic factors in the clearance of *ccna1* and, to a lesser extend, *nanog* mRNAs

## Methods

### Data Availability

Raw reads that support the findings of this study will be publicly available in the Sequence Read Archive under SRP114782 upon acceptance in a peer-reviewed journal. Meanwhile, we will be glad to share our raw data with anyone upon request.

### Zebrafish maintenance

Wild-type zebrafish embryos were obtained through natural mating of TU-AB strain of mixed ages (5-18 months). Mating pairs were randomly chosen from a pool of 60 males and 60 females allocated for each day of the month. Fish lines were maintained following the International Association for Assessment and Accreditation of Laboratory Animal Care research guidelines, and approved by the Yale University Institutional Animal Care and Use Committee (IACUC).

### *In vivo* and *in vitro* DMS modification

For *in vivo* DMS modification, 150 embryos at the specified stage were transferred to 5 mL eppendorf tubes containing 400 μL of system water from the fish facility. 100% DMS (Sigma-Aldrich) was diluted in 100% ethanol to obtain a 20% DMS stock solution. The DMS stock solution was used to generate a master mix containing 6% DMS and 600 mM Tris-HCl pH 7.4 (AmericanBio) in system water from the fish facility. The master mix solution was immediately mixed vigorously and 200 μL was added to each tube to reach a final concentration of 2% DMS and 200 mM Tris HCl pH 7.4. Embryos were incubated at room temperature for 10 min with occasional gentle mixing. The DMS solution was then quickly removed from the tubes and the embryos were flash frozen in liquid nitrogen. Frozen embryos were thawed and actively lysed with 800 μL of TRIzol (Life Technologies) supplemented with 0.7 M β-mercaptoethanol (Sigma-Aldrich) to quench any remaining trace of DMS. After 2 min incubation, TRIzol was added to reach a final volume of 4 mL and total RNA extracted following the manufacturer’s protocol. Poly(A)+ mRNAs were isolated using oligo d(T)_25_ magnetic beads (New England BioLabs) following the manufacturer’s protocol and eluted in 20 μL of water.

For *in vitro* DMS modification, embryos were collected and total RNA extracted as mentioned above, omitting DMS and β-mercaptoethanol. For each replicate, 20 μg of total RNA was resuspended in RNA folding buffer (57 mM Tris HCl pH 7.4 and 114 mM KCl) in a final volume of 44 μL. RNA samples were incubated at 65 °C for 2 min and slowly brought back to 28 °C over ∼45 min in a heating block. Then, 5 μL of 100 mM MgCl_2_ was added to each tube and samples were incubated at 28 °C for 3 min. RNA was modified by adding 1 μL of 25% DMS, previously diluted in ethanol (1:3 dilution), to reach a final DMS concentration of 0.5%. The reaction was incubated at 28 °C for 10 min in a PCR machine and quenched by the addition of 59 μL of stop solution (3 M β-mercaptoethanol, 508 mM sodium acetate and 15 μg glycoblue). After a 5 min incubation at room temperature, modified RNA samples were ethanol precipitated by the addition of 220 μL ethanol. Poly(A) + mRNAs were isolated as mentioned above.

All DMS-seq experiments (*in vivo, in vitro*, untreated, PatA-treated, and CHX-treated at 2 hpf, as well as 4 and 6 hpf) were performed in biological duplicates from embryos originated from different crosses and collected on different days.

### Translation inhibitor treatments

PatA reduces the levels of functional eIF4F initiation complex, which is essential for cap-dependent translation^51^, by trapping eIF4A onto mRNAs and ectopically enhancing its RNA helicase activity^36,37^ (Fig. 3a). Cycloheximide (CHX) binds to the E-site of the 60S ribosome subunit, inhibiting ribosomal translocation during translation elongation^38^. For DMS-seq and ribosome profiling experiments, 16-cell stage embryos were bathed in either 10 μM pateamine A (PatA) or 50 μg/ mL cycloheximide (CHX). Embryos were collected once untreated embryos from the same clutches reached 64-cell stage (2 hpf) (∼45 min following the addition of inhibitors). For DMS-seq samples, 150 embryos were collected in a 5 mL tube for each condition (untreated, PatA and CHX) and DMS-treated. Translation inhibitor concentrations were maintained during the DMS modification step. For ribosome profiling samples, 55 embryos per-condition and per-replicate were collected and flash frozen in liquid nitrogen.

### DMS-seq library preparation

DMS-seq experiments were performed as previously described by Rouskin et *al*^13^, with minor changes. Briefly, DMS treated poly(A)+ RNA samples were denatured at 95 °C for 2 min and fragmented at 95 °C for 1.5 min in 1X RNA fragmentation buffer (Zn^2+^ based from Ambion). The reaction was stopped by adding 0.1 volume of a 10X Stop solution (Ambion). One volume of formamide loading dye (95% formamide, 10 mM EDTA, 0.025% bromophenol blue and 0.025% xylene cyanol) was added to fragmented RNAs, and samples were separated in a 10% TBU (Tris borate 8 M urea) PAGE. Fragments were visualized by blue light (Clare Chemical Research), and RNAs of 60-70 nucleotides were excised. Gel pieces were shredded and gel extracted in 300 μL of nuclease free water (AmericanBio) at 70 °C for 10 min with vigorous shaking. Eluted RNA was purified using Spin-X tube filters (Sigma-Aldrich) and ethanol precipitated by adding 33 μL 3 M sodium acetate, 1.5 μL glycoblue and 700 μL ethanol. RNA samples were dissolved in 6 μL of nuclease-free water, and 3’ phosphates were removed by incubating the samples at 37 °C for 1 h with 1 μL 10X PNK buffer (NEB), 1 μL SUPERase Inhibitor (Ambion) and 2 μL of T4 PNK enzyme (NEB). Samples were then directly ligated to 1 μg of our in-house barcoded 3’ adapters, either /5rApp/ NNcacaACTGTAGGCACCATCAAT/3ddC/ or /5rApp/ NNagagACTGTAGGCACCATCAAT/3ddC/ (IDT DNA) —in-house barcode depicted in lower case— by adding 1 μL 0.1 M DTT, 6 μL 50% PEG, 1 μL 10X ligase2 buffer, 2 μL T4 RNA ligase2, truncated K227Q (NEB), and incubated at 25 °C for 1.5 h. Each replicate was ligated to a different barcode set, and therefore, replicates could be pooled following ligation. 20 μL of formamide dye was added to each tube, and ligated products were run in a 10% TBU PAGE for ∼45 min, visualized by blue light and separated from unligated adapters by gel extraction as described above. Reverse transcription was performed in 20 μL at 52 °C for 45 min using Superscript III (Invitrogen), 5 ’-(Phosphate)-NNNNAGATCGGAAGAGCGTCGTGTAGGGAAAGAGTGTAGATCTCGGTGGTCGC-(SpC18)-CACTCA-(SpC 18) – TTCAGACGTGTGCTCTTCCGATCTATTGATGGTGCCTACAG-3’ (IDT DNA) reverse transcription primer, followed by RNAse H treatment at 37 °C for 15 min. cDNAs were separated in a 10% TBU PAGE for 1.5 h and truncated reverse transcription products of 25-45 nucleotides above the size of the reverse transcription primer were extracted by gel purification. Samples were then circularized using CircLigase (Epicentre), ethanol precipitated and PCR amplified using Illumina sequencing adapters, keeping the number of cycles to the minimum needed for the detection of amplified products (9-11 cycles).

### Tetrahymena ribozyme spike-in control

Tetrahymena ribozyme sequence was ordered from IDT DNA. RNA was generated using a AmpliScribe-T7-Flash transcription kit (Epicentre) and *in vitro* DMS-probed as described above. DMS-modified *Tetrahymena* ribozyme was then spiked in to untreated poly(A)+ RNA sample extracted from 2 hpf embryos, random-fragmented and subjected to the DMS-seq library preparation protocol described above.

### Ribosome profiling experiments

Ribosome profiling was performed using the Artseq Ribosome Profiling Kit Mammalian (Epicentre) as previously described^52^ with minor changes. For each condition, two biological replicates were generated, originated from embryos using different crosses and collected at different days. For each replicate, 55 embryos were lysed in 800 μL of lysis buffer (1X polysome buffer, 1% Triton X-100, 1 mM DTT, 25 U/mL DNaseI and 100 μg/mL cycloheximide) according to the manufacturer’s protocol (Epicentre). Lysates were centrifuged for 10 min at 20,000 g at 4 °C. 4.5 μL of ARTseq Nuclease was added to 600 μL of lysate supernatant and incubated at 25 °C for 45 min with gentle mixing. Nuclease digestion was stopped by addition of 22.5 μL of SUPERase-In^TM^ RNase Inhibitor (Life Technologies) and chilled on ice for 5 min. Ribosomes were purified by Sephacryl S400 spin column chromatography (GE Healthcare) according to manufacturer’s protocol. Prior to RNA purification, 7.5 μL of a mix of four different 28 nucleotide spike-in RNA oligos (3.5 × 10^−10^ molar 5’-NAAGTCTAGKACTAGTAAMTGATCATAN-3’, 6.9 × 10 ^− 11^ molar 5 ’ - NACTAGTAGKAGATCATAMTTCATAGAN-3’, 1.4 × 10 ^− 11^ molar 5 ’ - NAGATCATGKATCATAGAMTAGTCTAAN-3’, 2.8 × 10 ^− 12^ molar 5 ’ - NATCATAGGKAAGTCTAAMTCTAGTAAN-3’, where N is any of the 4 nucleotides, K is either G or T and M either A or C) was added to the pooled purified ribosome samples. Purified ribosome protected fragments were separated in a 15% TBU gel and fragments of 28 to 30 nucleotides were extracted. Adapters were ligated at the 3’-end of the fragments, reverse transcribed, gel purified and circularized as described above for DMS-seq fragments.

For RNA-seq samples (input), 75 ng of *Saccharomyces cerevisiae* RNA was added as spike-in to 175 μL of the remaining lysate supernatant, followed by extraction of total RNA using TRIzol. Total RNA samples were sent to the Yale Center for Genome Analysis and strand-specific TruSeq Illumina RNA sequencing libraries were constructed. Prior to sequencing, samples were treated with Epicentre Ribo-Zero Gold according to the manufacturer’s protocol.

### RNA-seq time course experiments

To quantify changes in mRNA abundance and to measure the impact of zygotic factors and miR-430 activity, we performed an RNA-seq time course experiment in wild type conditions, in the presence of alpha-amanitin (to inhibit zygotic transcription activation), and in the presence of an miR-430 inhibitor. To inhibit transcription activation of the zygotic genome, embryos were injected with 2 ng of alpha-amanitin (aAm) (Sigma Aldrich), which is an inhibitor of RNA polymerase II. To inhibit miR-430 activity, embryos were injected with 1 nanoliter of 10 μM of tiny locked nucleic acid (tinyLNA), complementary to the seed region of miR-430 (5’-TAGCACTT-3’ (Exiqon)). ∼25 embryos were collected per-stage and per-condition. Embryos were lysed in TRIzol and spiked with 75 ng of *Saccharomyces cerevisiae* RNA for normalization purposes. Total RNA was extracted, sent to the Yale Center for Genome Analysis and strand-specific TruSeq Illumina RNA sequencing libraries were constructed for each sample. Prior to sequencing, samples were treated with either Epicentre Ribo-Zero Gold, to deplete ribosomal RNA, or oligo dT beads, to enrich in poly(A)+ RNA, according to manufacturer’s protocol.

### Read sequencing and mapping

DMS-seq, ribosome profiling and RNA-seq samples were sequenced on Illumina HiSeq 2000/2500 machines producing single-end 76 nucleotide reads. This yielded over 1 billion reads, 200 million reads and 100 million reads for DMS-seq, ribosome profiling, and RNA-seq respectively. Sequencing samples are summarized in Extended Data Table 1.

Following the library preparation protocol for DMS-seq and ribosome profiling samples, raw reads contained the following features: NNNN-insert-NN-barcode(4-mer)-adapter where the 6N (NNNN+NN) sequence composes the Unique Molecular Identifier (UMI), “barcode “is the sample 4-mer in-house barcode and adapter is the 3’-Illumina adapter. The UMI was used to discard PCR duplicates and count single ligation events. The barcode was used to mark individual replicate following the 3’-adapter ligation step. Basecalling was performed using CASAVA-1.8.2. The Illumina TruSeq index adapter sequence was then trimmed by aligning its sequence, requiring a 100% match of the first five base pairs and a minimum global alignment score of 60 (Matches: 5, Mismatches: −4, Gap opening: −7, Gap extension: −7, Cost-free ends gaps). Trimmed reads were demultiplexed based on the sample’s in-house barcode, the UMI was clipped from the 5’- and 3’-ends and kept within the read name, for marking PCR duplicates. Reads were then depleted of rRNA, tRNA, snRNA, snoRNA and miscRNA, using Ensembl78 annotations^53^, as well as from RepeatMasker annotations, using strand-specific alignment with Bowtie2 v2.2.4^54^. The remaining reads were aligned to the zebrafish Zv9 genome assembly using STAR version 2.4.2a^55^ with the following non-default parameters: *- - align Ends Type End To End - - outFilterMultimapNmax 100 –seedSearchStartLmax 15 --sfbdScore 10 --outSAMattributes All*. Genomic sequence indices for STAR were built including exon-junction coordinates from Ensembl 78. Only reads of unique UMI were kept at each genomic coordinate for DMS-seq and ribosome profiling experiments. For ribosome profiling samples, reads were also aligned to the 256 sequences composing the 4 partially degenerated 28 nucleotides RNA spike-ins using Bowtie2 v2.2.4.

Raw reads from RNA-seq experiments were processed using the same pipeline, omitting the adapter trimming, barcoding demultiplexing and UMI clipping steps. The filtered reads were aligned onto Zebrafish Zv9 and *Saccharomyces cerevisiae* r64d1d1 genome assemblies using STAR, with the same parameters as described above. For both species, STAR genomic sequence indices were built including exon-junction coordinates from Ensembl 78.

### DMS-seq profiles and accessibilities

Per-transcript profiles were computed using uniquely mapped reads overlapping at least 10 nucleotides with the transcripts’ annotation. Each read count was attributed to the nucleotide in position –1 of the read’s 5’-end within the transcript coordinate, to correct for the fact that reverse transcription stops one nucleotide prior to the DMS-modified nucleotide. To determine read distributions for each nucleotide (Extended Data Fig. 1c), only transcripts with a minimum of 100 counts were considered. In the same figure, ‘Transcriptome’ represents the frequency of each nucleotide for the same subset of transcripts (Extended Data Fig. 1c).

Accessibilities were calculated following the 2%/8% rule^56^, i.e., by normalizing the read counts proportionally to the most reactive A and C bases within the region after the removal of outliers. More specifically, the 2% most reactive A and C bases were discarded and each position was divided by the average of the next 8% most reactive A and C bases. Accessibilities greater than 1 were set to 1, and accessibilities for G and T were set to 0.

### Calculating translation efficiency

Translation efficiency was calculated by dividing the Ribo-seq RPKM by the RNA-seq RPKM (RPKM; Read Per Kb and per Million reads of spike-in) for each coding sequence, excluding the first and last three codons (effective CDS) as described in Bazzini *et al.*^52^. Using the effective CDS of each transcript allows computation of translation efficiency from actively translating ribosomes.

For ribosome profiling samples, the effective CDS annotation was shifted 12 nucleotides upstream to position each read at the ribosome P-site location. RPKMs were computed by summing the effective CDS counts, including reads matching up to five times in the genome (each mapping site counting 1/n, n = number of mapping sites), and normalizing with the effective CDS length and the total number of reads mapped to the spike-in. For RNA-seq, RPKMs were calculated using the same pipeline applied to obtain ribosome profiling RPKMs, with the following differences: the effective CDS annotation remained unshifted, counts from reads overlapping the effective CDS by a minimum of 10 nucleotides were added, and the total number of reads mapped to yeast RNAs was used as normalization spike-in. Finally, translation efficiency was computed by dividing ribosome profiling RPKMs by RNA-seq RPKMs for each transcript effective CDS. For each sample, this analysis was performed by combining reads from both replicates.

### Identification of conserved RNA structures

Conserved RNA structures (Extended Data Fig. 1g, h) were identified using the RNAz software^57^ on the UCSC *D. rerio* multiz 8-way whole genome alignment (http://hgdownload.cse.ucsc.edu/goldenPath/danRer7/multiz8way), with default parameters.

### mRNA accessibility analysis

To investigate global mRNA structure, the average accessibility of 5’-UTR, CDS and 3’-UTR was individually calculated and compared across conditions and/or between translation statuses. For this aim, we first identified the major transcript isoform of each gene, by selecting the one with the highest DMS-seq read density and with UTRs longer than 75 nucleotides (total of 11,404 mRNAs). If no isoform met the minimum UTR length criteria, no isoform was selected for that gene. Transcripts with average DMS-seq count per A/C equal to or greater than 5 (except when specified differently), and with counts covering at least 85% of A/C in each region (UTRs and CDS) were selected for further analysis. When the analysis was performed on full-length UTRs (Fig. 4c and Extended Data Fig. 1l, 2b, 2c, 3a, 5a and 6a), DMS-seq data was used to determine the final UTR lengths. Specifically, UTR length was extended one nucleotide at a time, starting at 76, until the A/C counts coverage fell under 85%. The longest 5’- and 3’-UTRs with at least 85% coverage was then associated to this transcript. Accessibilities were calculated within each region (5’-UTR, CDS and 3’-UTR) of each transcript as described above and averaged (for A/C bases only) to obtain a single value per region. For all global accessibility mRNA region analyses (cumulative plots), statistical significance between highly (red) and lowly (blue) translated mRNAs within the same condition was calculated using a one-sided Mann-Whitney *U* test while statistical significance between the same subset of mRNAs, but across two different conditions, was calculated using a two-sided Wilcoxon signed-rank test.

### Definition and selection of differentially translated mRNAs

To select lowly and highly translated mRNAs, transcripts were binned in quintiles based on translation efficiency. The quintile with the lowest translation efficiency was labeled as ‘low translation’ and the one with the highest translation efficiency was labeled as ‘high translation’.

### Sliding windows analysis

To decipher which regions of the transcript were significantly changing in RNA structure across conditions and developmental stages, per-transcript DMS-seq counts were subdivided into overlapping 100-nt sliding windows, offset by one nucleotide. Only windows with minimum coverage of 250 counts were considered for further analysis. Windows with statistically significant changes in RNA structure across conditions were identified using the Kolmogorov-Smirnov test, a non-parametric test sensitive to differences both in location and shape of the empirical cumulative distribution functions of two samples. The Gini index, which has been previously used to compare RNA structures^13^ due to its ability to capture inequalities in a given distribution, was used to identify windows with increased or decreased RNA structures (Fig. 1c). All analyses were performed separately for each replicate, and *P*-values and Gini indices were combined using Fisher’s method (R package ‘metap’) and averaged, respectively. Differential structured windows between two conditions were defined as those that were significant (combined *P*-value<0.05 in the KS-test, and that had an average Gini fold change across replicates greater than 1.1 or smaller than 0.9 (Fig. 1c). Metaplots of the distribution of differentially structured windows along the transcript were normalized by the region lengths and by the total number of windows analyzed in each region (5’-UTR, CDS, 3’-UTR).

### Sequence conservation analysis

To determine the sequence conservation, we employed PhastCons regional conservation^58^ scores derived from the Multiz^59^ alignment of 8 vertebrate species, which included 5 fish species (ftp://hgdownload.cse.ucsc.edu/goldenPath/danRer7/phastCons8way/).

### uORF analysis

A single transcript isoform per gene was selected as described above except for the minimal UTR length, which was set to 12 nucleotides. We then used the zebrafish uORFs dataset from Johnstone *et al.*^60^, which contains location and translation efficiency information, to identify lowly and highly translated uORFs at 2 hpf as described above for mRNAs. Per-uORF DMS-seq accessibility profiles were calculated by applying the 2%/8% rules mentioned above, for uORFs longer than 150 nucleotides. For shorter uORFs, DMS accessibilities were computed using 150 nucleotide regions starting at the uORF AUG initiation codon. Global uORF accessibilities were derived as described above. uORFs containing less than 10 A/C bases were removed from this analysis.

### Poly(A) tail length and miR-430-mediated repression analyses

Poly(A) tail length datasets were obtained from Subtelny et *al.*^33^, who developed a technique, called poly(A)-tail length profiling by sequencing (PAL-seq), to measure the poly(A) tail length of transcripts from various organisms, including 2, 4 and 6 hpf zebrafish embryos. Only transcripts with at least 50 poly(A) tags were considered in our poly(A) tail length analyses (Fig. 4b, c and Extended Data Fig. 5). To analyze the impact of miR-430-mediated repression on mRNA structures (Fig. 4d, e), a set of 483 mRNAs targeted by miR-430 was selected using the following criteria: i) minimum 2-fold decrease in mRNA level between 2 and 6 hpf (6/2); ii) minimum 1.5-fold increase in mRNA level between 6 hpf tinyLNA and 6 hpf wild type (6tinyLNA/6wt); and iii) minimum of 5 reads on average per A/C base in DMS-seq experiments. For comparison, a random set of 483 mRNAs was selected among all the remaining transcripts with enough DMS-seq coverage. For genes with multiple transcript isoforms, the one with UTRs greater than 50-nt in length and with the highest coverage in our DMS-seq experiment at 2 hpf was selected. If no transcript isoform met those criteria for a specific gene, no transcript from that gene was included in the analysis.

### RNA secondary structure analyses

Predicted RNA secondary structures were computed using either the *Fold* executable of the RNA structure package^61^ (version 5.6) or the SeqFold software^40^. For the *Fold* software, the default parameters were used except for the temperature parameter, which was set to 301.15 Kelvin (28 °C)^62^. DMS-seq accessibilities of A and C bases were inferred as soft constraints using the – dms option. The Gibbs energy of folding (ΔG) of each predicted RNA structure was calculated using the *efn2* executable with default parameters except for the temperature parameter, which was set to 301.15 Kelvin (28 °C)^63^. Arc plots representing RNA secondary structure were drawn using the R-chie software with default parameters^64^. The analysis of the RNA structure surrounding AUG initiation codons was performed using 100-nt windows centered on each AUG with a minimum DMS-seq depth of 5 reads in average per A and C bases (Fig. 2e and Extended Data Fig. 2d-g). The analysis of the impact of ribosomes on local RNA structure was limited to 100-nt windows located in CDS regions and with a minimum DMS-seq depth of 5 reads in average per A and C bases (Fig. 3f and Extended Data Fig. 3b, c). Windows with ribosome footprints correspond to those with an average of ribo-seq reads between replicates greater than 7. Windows without ribosome footprints correspond to those with an average of ribo-seq reads lower than 1 and found in mRNAs with translation efficiency lower than 1.65. Each subset contains a total of 728 non-overlapping windows covering 332 and 312 different genes for windows with and without ribosome footprints, respectively (Extended Data Fig. 3c).

For the SeqFold analysis, only transcripts with more than 7 reads in average per A/C bases in all four tested conditions (untreated, *in vitro*, PatA-treated or CHX-treated) were considered, which resulted in a set of 1,143 transcripts. DMS-seq accessibility of A and C bases was inferred following the same protocol as for SHAPE accessibilities and energies of folding were directly retrieved from the SeqFold output. To determine the global similarity of predicted RNA structures across conditions, per-transcript Gini indices were computed from the SeqFold outputs using the reldist library in R. Principal Component Analysis (PCA) was then used to reduce the dimensionality of the dataset. A biplot of the PCA loadings (i.e., conditions/treatments) of the two first principal components was used to determine which conditions were most similar. The same transcriptome-wide PCA analysis was done separately for CDS and 3’-UTR regions. The cumulative proportion of variance explained by the two first components was: 92% (PC1) and 3.1% (PC2) in CDS regions, and 94.5% (PC1) and 2.3% (PC2) in 3’-UTR regions.

### miR-430 site structural analysis

To assess the impact of RNA structures on microRNA activity, we selected 18 different endogenous miR-430 binding sites with DMS-seq coverage equal to or greater than 5 reads per A/C bases and obtained their decay activity from Yartseva et *al.*^45^. The stability (minimum free energy, ΔG) of the structure formed by the 120-nt region centered on the miR-430 seed was calculated in different conditions (*in silico, in vitro* and *in vivo* at 2 hpf, prior to miR-430 expression), and the correlation between the RNA structure stability and decay strength was determined using Spearman correlations (Fig. 5d). The predicted secondary structure and stability of a 200-nt region centered on the miR-430 seed were computed for additional endogenous miR-430 sites found in maternal mRNA 3’-UTRs (Fig. 5e, i and Extended Data Fig. 8). For each maternal mRNA containing miR-430 sites, decay patterns were analyzed in the wild type condition (with miR-430, WT) or when miR-430 activity was inhibited (-miR-430) by using a tiny LNA complementary to the seed region of miR-430. An increase in maternal mRNA stability in the –miR-430 condition implies miR-430-mediated repression of those maternal mRNAs in WT condition (Fig. 5f, j and Extended Data Fig. 8).

Next, GFP reporters containing endogenous miR-430 binding sites in their 3’-UTR were used, and GFP expression was quantified in both wild type and MZ*mir430* mutant fish, where the miR-430 locus on chromosome 4 has been deleted^65^ (Fig. 5g, h, k, l). In addition to the 200-nt wild type sequence of each endogenous miR-430 site, mutated versions that either stabilize or destabilize the *in vivo* structure were also built. Briefly, the GFP coding sequence and miR-430 sites were cloned in pCS2 plasmid between the BamHI and XhoI sites. The resulting plasmids were linearized with NotI digestions, and the products were *in vitro* transcribed with the Sp6 mMessage mMachine kit (Thermo Fisher, cat. No. AM1340). Zebrafish embryos were injected with 100 pg of mRNA GFP reporters and 100 pg of DsRed reporter (internal control) at the 1-cell stage. Fluorescence was quantified at 24 hpf using ImageJ. MiR-430 activity was calculated as the ratio GFP/DsRed for both genetic backgrounds (wild type and MZ*mir430*). Wild type GFP/DsRed was then normalized by the GFP/DsRed ratio from MZ*mir430*, obtaining miR-430-mediated repression values for each reporter. Reported values come from three biological replicates (Fig. 5h, l).

Finally, to determine the impact of Ago2/ miR-430 complex binding on the RNA accessibility of the recognition site, we analyzed the per-nucleotide accessibilities of 100-nt windows centered on individual miR-430 seeds (8-and 7-mers) found in the 3’-UTR of miR-430 targets (identified as described above), for each developmental stage and condition (Extended Data Fig. 7e-g). For comparison, per-nucleotide accessibilities of randomly chosen 100-nt windows within the same set of 3’-UTRs were used. Global seed and control accessibilities were calculated by averaging the accessibility of each A and C bases found in the 8-nt sequence located in the center of each window (Extended Data Fig. 7f). Only windows with a minimum of 5 reads in average per A and C bases were considered.

### RNA-element selection assay

To quantify the regulatory activity of maternal mRNA 3’-UTR sequences during the MZT, we used an RNA-element selection assay (RESA) as described in Yartseva *et al.*^45^ with minor changes (Fig. 6b). Maternal 3’-UTR regions of 200 nucleotides showing changes in RNA structure during the MZT (based on KS-test across stages, *P*<0.05) or no change (as controls, *P*>0.05) were amplified from cDNA of 2 hpf embryos (see Extended Data Table 2 for primer sequences). In total, 74 dynamic and 25 control regions were amplified and inserted in the 3’-UTR of a GFP reporter by assembly PCR with Phusion High Fidelity DNA polymerase (NEB, cat. No. M0530), and purified on agarose gel. The final library constructs consisted of an Sp6-promoter, a GFP coding sequence, a 3’-UTR with various inserts flanked by part of the IIlumina 5 ’- and 3 ’-adaptor sequences, and a SV40 polyadenylation signal. The library was *in vitro* transcribed with an Sp6 mMessage mMachine kit (Thermo Fisher, cat. No. AM1340) and the resulting RNA library was injected in 1-cell stage embryos, with or without the presence of alpha-amanitin. Alpha-amanitin (aAm) is an inhibitor of RNA polymerase II that blocks the activation of the zygotic genome. Approximately 25 injected embryos were collected at 2 and 6 hpf (with and without aAm), and total RNA was extracted with TRIzol reagent according to the manufacturer’s protocol. Poly(A)+ mRNA was selected with NEB oligo d(T)_25_ magnetic beads (NEB, cat. no. S1419S) following the manufacturer’s protocol. Poly(A)+ mRNA was recovered with ethanol precipitation and dissolved in 11 μL of water. Concentrated poly(A)+ m RNA was reverse-transcribed with a reporter specific primer (5’-CATCAATGTATCTTATCATGTCTGGATC-3’) with SuperScript III. Illumina adapters were added and the library amplified with ∼20 circles of Phusion PCR with the forward primer (5 ’-AATGATACGGCGACCACCGAGATCTACACTCTTTCCCTACACGACGCTCTTCC-3’) and the reverse primer (5 ’-CAAGCAGAAGACGGCATACGAGAT(barcode)GTGA CTGGAGTTCAGACGTGTGCTCTTCCGATCT-5’). The RESA experiment was performed in five biological replicates.

RESA libraries were sequenced with Illumina HiSeq 2000/2500 machines producing single-end 76-nt reads. Reads were processed and aligned as mentioned above for DMS-seq reads, omitting the adapter trimming, barcoding demultiplexing and UMI clipping steps. The number of reads corresponding to each insert was counted and normalized by the total number of reads in each sample. Changes in reporter abundance during the MZT were calculated as the ratio of counts (6/2 hpf or 6wt/6aAm) for each insert, for those reporters with enough coverage at 2 hpf (average count greater or equal to 250). Then, the average of the replicates was calculated after removing outliers (lower/higher of the mean ± standard deviation) for each insert. When multiple inserts corresponding to dynamic structure or control were present in a single 3’-UTR, the insert with the highest impact on gene expression (positive or negative) was kept for each category. This analysis allowed quantifying the impact of 53 and 18 3’-UTR sequences with dynamic structures and with no structural change, respectively, in regulating the mRNA abundance during the MZT (Fig. 6c, d, f, h and Extended Data Fig. 9b, c, e).

### KHSRP iCLIP experiment and analysis

To identify the regions bound by KHSRP during early embryogenesis, we performed iCLIP experiment as described in Huppertz *et al.*^66^ with minor changes. Embryos at sphere stage (4 hpf) were collected and irradiated with 254 nm UV light to induce crosslinking (embryos were snap frozen and stored in batches to yield a total of 1,000 embryos/condition). Frozen embryos were then thawed and homogenized on ice in iCLIP lysis buffer^66^. An affinity-purified rabbit polyclonal antibody raised against zebrafish KHSRP (generated by YenZym Antibodies, LLC) was used to isolate RNA-protein complexes. Briefly, 200 μl Protein G Dynabeads were added to 50 μg antibody in lysis buffer. Beads were incubated for an hour, washed three times with lysis buffer and added to the lysates. As a control, a parallel experiment was performed without the addition of antibody. Subsequent steps in the iCLIP protocol were performed as described in Huppertz *et al*.^66^, except for the 3’-adaptors containing in-house barcode, the reverse transcription primer and PCR primers, which were identical to the ones used for the DMS-seq library preparation. Twenty cycles of amplification of cDNA were used for final library construction. Barcoded PCR-amplified libraries were size-selected in a 6% TBE gel and combined for Illumina sequencing. Both the KHSRP pull-down and the no-antibody control were performed in triplicates. Raw reads from iCLIP experiments were processed using the same pipeline than for the DMS-seq experiments.

To define the binding preference of KHSRP, bound windows were defined for each 6-mer with at least 10 reads and extended 5 nucleotides upstream and 10 nucleotides downstream. For each bound window, the frequency of each 6-mer was computed. These frequencies were normalized by the average 6-mer frequencies observed in the no-antibody controls, which were calculated as mentioned above except for the read depth requirement, which was set to 2 instead of 10 reads to compensate for differences in sequencing depth. To build the logo representation, the top 10 most frequent normalized 6-mers was aligned using MAFFT with the following parameters: *--reorder --lop -10 --lexp -10 --localpair --genafpair --maxiterate 1000*. Multiple sequence alignment positions with more than 50% gaps were trimmed on both ends. Nucleotide frequencies were calculated using the final trimmed alignment, and represented as logo (Extended Data Fig. 7a).

iCLIP peaks were called by testing scanning 30-nt windows using Poisson’s law. Overlapping significant windows (*P*<0.01) were then merged to obtain the peaks. To get robust Poisson parameter estimates, only transcripts with a minimum of 20 positions with at least 1 read were analyzed. Final KHSRP peaks correspond to overlaps of peaks found in at least 2 out of 3 replicates. To analyze the RNA structure of KHSRP-bound and -unbound regions, we defined bound regions as 100-nt windows centered on the middle point of each KHSRP peak. For each bound region, an unbound control within the same 3’-UTR and overlapping with less than 25% with the bound region was selected. The accessibility of each region was computed by averaging the accessibility of A and C bases. Finally, only transcripts and regions with a minimum of 5 reads in average per A and C bases were considered (Extended Data Fig. 7c).

## References

1. Sharp, P. A. The centrality of RNA. Cell 136, 577–580 (2009).

2. Warf, M. B. & Berglund, J. A. Role of RNA structure in regulating pre-mRNA splicing. Trends Biochem. Sci. 35, 169–178 (2010).

3. Martin, K. C. & Ephrussi, A. mRNA localization: gene expression in the spatial dimension. Cell 136, 719–730 (2009).

4. Ray, P. S. et al. A stress-responsive RNA switch regulates VEGFA expression. Nature 457, 915–919 (2009).

5. Weingarten-Gabbay, S. et al. Systematic discovery of cap-independent translation sequences in human and viral genomes. Science 351, aad4939–aad4939 (2016).

6. Kedde, M. et al. A Pumilio-induced RNA structure switch in p27-3’ UTR controls miR-221 and miR-222 accessibility. Nat. Cell Biol. 12, 1014–1020 (2010).

7. Lucks, J. B. et al. Multiplexed RNA structure characterization with selective 2’-hydroxyl acylation analyzed by primer extension sequencing (SHAPE-Seq). Proc. Natl. Acad. Sci. U.S.A. 108, 11063–11068 (2011).

8. Kwok, C. K., Tang, Y., Assmann, S. M. & Bevilacqua, P. C. The RNA structurome: transcriptome-wide structure probing with next-generation sequencing. Trends Biochem. Sci. 40, 221–232 (2015).

9. Mortimer, S. A., Kidwell, M. A. & Doudna, J. A. Insights into RNA structure and function from genome-wide studies. Nat. Rev. Genet. 15, 469–479 (2014).

10. Ding, Y. et al. In vivo genome-wide profiling of RNA secondary structure reveals novel regulatory features. Nature 505, 696–700 (2014).

11. Kertesz, M. et al. Genome-wide measurement of RNA secondary structure in yeast. Nature 467, 103–107 (2010).

12. Lu, Z. et al. RNA Duplex Map in Living Cells Reveals Higher-Order Transcriptome Structure. Cell (2016). doi:10.1016/j.cell. 2016.04.028

13. Rouskin, S., Zubradt, M., Washietl, S., Kellis, M. & Weissman, J. S. Genome-wide probing of RNA structure reveals active unfolding of mRNA structures in vivo. Nature 505, 701–705 (2014).

14. Spitale, R. C. et al. Structural imprints in vivo decode RNA regulatory mechanisms. Nature 519, 486–490 (2015).

15. Guo, J. U. & Bartel, D. P. RNA G-quadruplexes are globally unfolded in eukaryotic cells and depleted in bacteria. Science 353, aaf5371 (2016).

16. Takyar, S., Hickerson, R. P. & Noller, H. F. mRNA helicase activity of the ribosome. Cell 120, 49–58 (2005).

17. Wen, J.-D. et al. Following translation by single ribosomes one codon at a time. Nature 452, 598–603 (2008).

18. Qu, X. et al. The ribosome uses two active mechanisms to unwind messenger RNA during translation. Nature 475, 118–121 (2011).

19. Burkhardt, D. H. et al. Operon mRNAs are organized into ORF-centric structures that predict translation efficiency. Elife 6, 811 (2017).

20. Kozak, M. Circumstances and mechanisms of inhibition of translation by secondary structure in eucaryotic mRNAs. Mol. Cell. Biol. 9, 5134–5142 (1989).

21. de Smit, M. H. & van Duin, J. Secondary structure of the ribosome binding site determines translational efficiency: a quantitative analysis. PNAS 87, 7668–7672 (1990).

22. Gray, N. K. & Hentze, M. W. Regulation of protein synthesis by mRNA structure. Mol Biol Rep 19, 195–200 (1994).

23. Dvir, S. et al. Deciphering the rules by which 5’-UTR sequences affect protein expression in yeast. PNAS 110, E2792–E2801 (2013).

24. Zur, H. & Tuller, T. Strong association between mRNA folding strength and protein abundance in S. cerevisiae. EMBO Rep. 13, 272–277 (2012).

25. Tadros, W. & Lipshitz, H. D. The maternal-to-zygotic transition: a play in two acts. Development 136, 3033–3042 (2009).

26. Yartseva, V. & Giraldez, A. J. The Maternal-to-Zygotic Transition During Vertebrate Development: A Model for Reprogramming. Curr. Top. Dev. Biol. 113, 191–232 (2015).

27. Lee, M. T., Bonneau, A. R. & Giraldez, A. J. Zygotic genome activation during the maternal-to-zygotic transition. Annu. Rev. Cell Dev. Biol. 30, 581–613 (2014).

28. Lee, M. T. et al. Nanog, Pou5f1 and SoxB1 activate zygotic gene expression during the maternal-to-zygotic transition. Nature 503, 360–364 (2013).

29. Leichsenring, M., Maes, J., Mössner, R., Driever, W. & Onichtchouk, D. Pou5f1 Transcription Factor Controls Zygotic Gene Activation In Vertebrates. Science 341, 1005–1009 (2013).

30. Heyn, P. et al. The Earliest Transcribed Zygotic Genes Are Short, Newly Evolved, and Different across Species. Cell Reports 6, 285–292 (2014).

31. Giraldez, A. J. et al. Zebrafish MiR-430 promotes deadenylation and clearance of maternal mRNAs. Science 312, 75–79 (2006).

32. Lund, E., Liu, M., Hartley, R. S., Sheets, M. D. & Dahlberg, J. E. Deadenylation of maternal mRNAs mediated by miR-427 in Xenopus laevis embryos. RNA 15, 2351–2363 (2009).

33. Subtelny, A. O., Eichhorn, S. W., Chen, G. R., Sive, H. & Bartel, D. P. Poly(A)-tail profiling reveals an embryonic switch in translational control. Nature 508, 66–71 (2014).

34. Bazzini, A. A. et al. Codon identity regulates mRNA stability and translation efficiency during the maternal-to-zygotic transition. The EMBO Journal 35, 2087–2103 (2016).

35. Li, F. et al. Regulatory Impact of RNA Secondary Structureacross the Arabidopsis Transcriptome. The Plant Cell Online 24, 4346–4359 (2012).

36. Low, W.-K. et al. Inhibition of Eukaryotic Translation Initiation by the Marine Natural Product Pateamine A. Mol. Cell 20, 709–722 (2005).

37. Bordeleau, M.-E. et al. RNA-Mediated Sequestration of the RNA Helicase eIF4A by Pateamine A Inhibits Translation Initiation. Chemistry & Biology 13, 1287–1295 (2006).

38. Schneider-Poetsch, T. et al. Inhibition of eukaryotic translation elongation by cycloheximide and lactimidomycin. Nat. Chem. Biol. 6, 209–217 (2010).

39. Kuzmenko, A. et al. Mitochondrial translation initiation machinery: Conservation and diversification. Biochimie 100, 132–140 (2014).

40. Ouyang, Z., Snyder, M. P. & Chang, H. Y. SeqFold: genome-scale reconstruction of RNA secondary structure integrating high-throughput sequencing data. Genome Res. 23, 377–387 (2013).

41. Lim, J., Lee, M., Son, A., Chang, H. & Kim, V. N. mTAIL-seq reveals dynamic poly(A) tail regulation in oocyte-to-embryo development. Genes Dev. 30, 1671–1682 (2016).

42. Bazzini, A. A., Lee, M. T. & Giraldez, A. J. Ribosome profiling shows that miR-430 reduces translation before causing mRNA decay in zebrafish. Science 336, 233–237 (2012).

43. Kertesz, M., Iovino, N., Unnerstall, U., Gaul, U. & Segal, E. The role of site accessibilityinmicro RNA targetrecognition. Nat. Genet. 39, 1278–1284 (2007).

44. Long, D. et al. Potent effect of target structure on microRNA function. Nature Structural & Molecular Biology 14, 287–294 (2007).

45. Yartseva, V., Takacs, C. M., Vejnar, C. E., Lee, M. T. & Giraldez, A. J. RESA identifies mRNA-regulatory sequences at high resolution. Nat. Methods 14, 201–207 (2017).

46. Nakano, S.-I., Miyoshi, D. & Sugimoto, N. Effects of Molecular Crowding on the Structures, Interactions, and Functions of Nucleic Acids. Chemical reviews 114, 2733–2758 (2013).

47. Tyrrell, J., McGinnis, J. L., Weeks, K. M. & Pielak, G. J. The Cellular Environment Stabilizes Adenine Riboswitch RNA Structure. Biochemistry 52, 8777–8785 (2013).

48. Wu, X. & Bartel, D. P. Widespread Influence of 3’ - End Structures on Mammalian mRNA Processing and Stability. Cell 169, 905–917.e11 (2017).

49. Yaman, I. et al. The Zipper Model of Translational Control. Cell 113, 519–531 (2003).

50. Halstead, J. M. et al. An RNA biosensor for imaging the first round of translation from single cells to living animals. Science 347, 1367–1671 (2015).

51. Anne-Claude Gingras, Brian Raught, A. & Sonenberg, N. eIF4 Initiation Factors: Effectors of m RNA Recruitment to Ribosomes and Regulators of Translation. http://dx.doi.org/10.1146/annurev.biochem.68.1.913 68, 913–963 (2003).

52. Bazzini, A. A. et al. Identification of small ORFs in vertebrates using ribosome footprinting and evolutionary conservation. The EMBO Journal 33, 981–993 (2014).

53. Yates, A. et al. Ensembl 2016. Nucleic Acids Res. 44, D710–6 (2016).

54. Langmead, B. & Salzberg, S. L. Fast gapped-read alignment with Bowtie 2. Nat. Methods 9, 357–359 (2012).

55. Dobin, A. et al. STAR: ultrafast universal RNA-seq aligner. Bioinformatics 29, 15–21 (2013).

56. Deigan, K. E., Li, T. W., Mathews, D. H. & Weeks, K. M. Accurate SHAPE-directed RNA structure determination. Proc. Natl. Acad. Sci. U.S.A. 106, 97–102 (2009).

57. Gruber, A. R., Findei, S., Washietl, S., Hofacker, I. L. & Stadler, P. F. RNAz 2.0: improved noncoding RNA detection. Pacific Symposium on Biocomputing (2010).

58. Siepel, A. et al. Evolutionarily conserved elements in vertebrate, insect, worm, and yeast genomes. Genome Res. 15, 1034–1050 (2005).

59. Blanchette, M. et al. Aligning Multiple Genomic Sequences With the Threaded Blockset Aligner. Genome Res. 14, 708–715 (2004).

60. Johnstone, T. G., Bazzini, A. A. & Giraldez, A. J. Upstream ORFs are prevalent translational repressors in vertebrates. The EMBO Journal 35, 706–723 (2016).

61. Reuter, J. S. & Mathews, D. H. RNAstructure: software for RNA secondary structure prediction and analysis. BMC Bioinformatics 11, 1 (2010).

62. Mathews, D. H. Using an RNA secondary structure partition function to determine confidence in base pairs predicted by free energy minimization. RNA 10, 1178–1190 (2004).

63. Mathews, D. H., Sabina, J., Zuker, M. & Turner, D. H. Expanded sequence dependence of thermodynamic parameters improves prediction of RNA secondary structure. J. Mol. Biol. 288, 911–940 (1999).

64. Lai, D., Proctor, J. R., Zhu, J. Y. A. & Meyer, I. M. R-chie: a web server and R package for visualizing RNA secondary structures. Nucleic Acids Res. 40, gks241–e95 (2012).

65. Liu, Y. et al. Inheritable and Precise Large Genomic Deletions of Non-Coding RNA Genes in Zebrafish Using TALENs. PLoS ONE 8, e76387 (2013).

66. Huppertz, I. et al. iCLIP: protein-RNA interactions at nucleotide resolution. Methods 65, 274–287 (2014).

67. Mathews, D. H. et al. Incorporating chemical modification constraints into a dynamic programming algorithm for prediction of RNA secondary structure. Proc. Natl. Acad. Sci. U.S.A. 101, 7287–7292 (2004).

68. Golden, B. L., Gooding, A. R., Podell, E. R. & Cech, T. R. A preorganized active site in the crystal structure of the Tetrahymena ribozyme. Science 282, 259–264 (1998).

69. Han, J. et al. Posttranscriptional Crossregulation between Drosha and DGCR8. Cell 136, 75–84 (2009).

70. Walter, N., Woodson, R. T. & Batey, R. T. Non-Protein Coding RNAs. (Springer Science & Business Media, 2008).

71. Tijerina, P., Mohr, S. & Russell, R. DMS footprinting of structured RNAs and RNA-protein complexes. Nat Protoc 2, 2608–2623 (2007).

